# Microbiome assembly and maintenance across the lifespan of bumble bee workers

**DOI:** 10.1101/2022.05.11.491538

**Authors:** Tobin J. Hammer, August Easton-Calabria, Nancy A. Moran

## Abstract

How a host’s microbiome changes over its lifespan can influence development and aging. As these temporal patterns have only been described in detail for humans and a handful of other hosts, an important next step is to compare microbiome dynamics across a broader array of host-microbe symbioses, and to investigate how and why they vary. Here we characterize the temporal dynamics and stability of the bumblebee worker gut microbiome. Bumblebees are a useful symbiosis model given their relatively well-understood life history and simple, host-specific gut bacterial communities. Furthermore, microbial dynamics may influence bumblebee health and pollination services. We combined high-temporal-resolution sampling with 16S rRNA gene sequencing, quantitative PCR, and shotgun metagenomics to characterize gut microbiomes over the adult lifespan of *Bombus impatiens* workers. To understand how hosts may control (or lose control of) the gut microbiome as they age, we also sequenced hindgut transcriptomes. We found that, at the community level, microbiome assembly is highly predictable and similar to patterns of primary succession observed in the human gut. At the same time, partitioning of strain-level bacterial variants among colonies suggests stochastic colonization events similar to those observed in flies and nematodes. We also find strong differences in temporal dynamics among symbiont species, suggesting ecological differences among microbiome members in colonization and persistence. Finally, we show that both the gut microbiome and host transcriptome—including expression of key immunity genes—stabilize, as opposed to senesce, with age. We suggest that in highly social groups such as bumblebees, maintenance of both microbiomes and immunity contribute to the inclusive fitness of workers, and thus remain under selection even in old age. Our findings provide a foundation for exploring the mechanisms and functional outcomes of bee microbiome succession, and for comparative analyses with other host-microbe symbioses.

## Introduction

Understanding how and why microbial communities change over time is a fundamental goal of microbial ecology (1–3). For host-associated microbiomes, the local environment can change dramatically as hosts grow, mature, and age, influencing their temporal dynamics (4–8). These dynamics may also have functional consequences, possibly influencing or regulating host development and life history processes (5, 9–11). We have an increasingly clear picture of microbiome succession in humans and in certain models for biomedical and symbiosis research [e.g., (12–18)], for which a range of methods have been used to describe dynamics of both microbes and host processes in great detail. Patterns can vary substantially across hosts. For example, in primary succession, stochastic colonization dynamics observed in *D. melanogaster* and *C. elegans* (13, 14, 19) contrast with predictable gut microbiome assembly in human infants and honeybees (20–22). On the other hand, convergent patterns are also observed, especially with respect to microbiome maintenance in old age. In humans, gut microbiome composition becomes more variable in the elderly, with losses of core symbiont species (23–26). In lab models, gut microbiomes also shift (though in various ways) in old age (17, 27–29); these shifts may constitute a form of senescence, both responding and contributing to deterioration of gut physiology and immunity (28, 30, 31). However, given major biological differences, it is difficult to explain why we see divergent or convergent successional trajectories among these groups.

Marker gene-based studies of microbiome succession from a greater diversity of hosts suggest a much broader array of temporal patterns in nature. For example, microbiome assembly can differ even between closely related hosts [e.g., humans and chimps (32)], and in some hosts, microbiome senescence does not seem to occur (33). However, typical marker gene (e.g., 16S rRNA) amplicon sequence datasets lack information on taxa not amplified by the chosen primer set, on absolute abundance, and on activity (e.g. live versus dead, replicating versus dormant). Short amplicon datasets of conserved genes also lack phylogenetic resolution below the species (i.e. ASV) level, masking subspecies or strain-level dynamics. Furthermore, many of these studies lack data on host processes that might impact microbes, such as immune responses, especially at spatial and temporal scales relevant to microbial dynamics. For a general understanding of how and why host-associated microbiomes change over time, it would be useful to develop a broader range of host-microbiome systems, studied with a comprehensive set of host-and microbe-level analyses.

Eusocial corbiculate bees (honeybees, bumblebees, and stingless bees) are a promising group for comparative, in-depth studies of microbiome dynamics. First, these bees are key pollinators in natural and agricultural ecosystems, and bacterial symbionts have functional roles in host health (34–36). Therefore, the dynamics of these microbes could have important consequences for bees as well as plants. Bees are threatened by a variety of anthropogenic stressors (37), and baseline temporal variability in the microbiome needs to be measured in order to observe perturbations and study resilience (38, 39). Second, these clades are related and are ecologically similar in many respects and share some conserved symbiont taxa, but they also differ in key life history and ecological traits, as well as in the composition and functional potential of their microbiomes (34, 35, 40). This contrast among close relatives provides an opportunity to study how host traits shape the evolution of microbiome dynamics. Third, social bees have host-specific and very simple gut microbiomes, dominated by just a few core bacterial lineages (41). This simplicity makes it easier to delve below community-level patterns to study the temporal dynamics of individual species and strains within the microbiome. As with plants (42), microbial taxa may exhibit different life history strategies that shape colonization and persistence. These strategies are poorly understood for microbes within host-associated communities, but are likely important for shaping community-level succession (43).

Gut microbiome temporal dynamics have been relatively well-studied in the Western honeybee (*Apis mellifera*). The gut microbiome is completely different between larvae and adult workers (44), and it continues to change in composition and abundance as workers age (15, 22, 45–47). In *A. mellifera*, however, adults go through a highly stereotyped sequence of tasks with age (48), such that age and task effects are difficult to disentangle. For example, honeybees do not defecate until they become foragers and leave the hive for the first time (49); this could contribute to a decrease in microbial abundance from young nurses to older workers (45). Furthermore, the oldest workers are those who overwinter and enter a phase with distinct metabolic, immunological, thermal, and behavioral (e.g., lack of defecation) characteristics (45, 50, 51). Given these unique age-related shifts, it is unclear to what extent gut microbial succession in *A. mellifera* will extend to other social bees.

Bumblebees (*Bombus* spp.) differ from honeybees in many ways that likely relate to microbiome dynamics (35). Workers exhibit comparatively weak age polyethism (temporal division of labor); tasks are generally carried out by workers of all ages, though some tasks are more likely to be performed at certain ages (52). Symbionts are transmitted between generations by a single queen, instead of by a large group of workers as in honeybees; this changes the bottleneck size, and potentially, selection on caste-specific maintenance processes (35). They also lack certain bacteria characteristic of honeybees and have gained *Candidatus* Schmidhempelia bombi (hereafter, *Schmidhempelia*) (41, 53). A unique practical advantage of bumblebees is that full colonies can be reared indoors. This provides an opportunity to study intrinsic aging processes under optimal conditions, in the absence of environmental variation, and to sample microbiomes of very old bees that would normally be rare due to extrinsic mortality. Moreover, the bumblebee gut microbiome seems uniquely prone to disturbance: field-collected workers are often found to lack the core symbionts and instead harbor opportunistic environmental bacteria (35, 54, 55). This phenomenon has been linked to colony age, but old colonies will also tend to have older workers on average (55). Whether microbiome disturbance is due to individual senescence has not been fully resolved.

Previous work has outlined the early stages of gut microbiome succession in bumblebees (56–58), but there is no information on what happens to the microbiome in old age. We also lack information on the temporal dynamics of endogenous processes (e.g., immunity) in the bee gut that control gut microbes, and may be controlled by them (59, 60). Gut physiology and immunity senesce in many animals [e.g., (8, 28, 30, 61)], but these processes—and senescence generally— may operate quite differently between different castes of eusocial insects (62, 63). In bumblebees, a solitary queen founds the colony and produces cohorts of (mostly) nonreproductive workers; reproductive offspring (queens and males) are produced toward the end of the colony cycle (64). Given that: *i*) age-specific survival probabilities are similar over much of the bumblebee worker lifespan (65, 66), *ii*) even old workers contribute to colony reproduction (52), and *iii*) there is a need to transmit the core gut symbionts—and not pathogens or parasites—to new queens (35), one may expect only minimal senescence of worker microbiomes, gut physiology, and immunity. Indeed, some aspects of systemic immunity in bumblebees remain stable or increase with age (67).

Our main research questions are: how is the bumblebee gut microbiome assembled and maintained through the lifespan—are patterns predictable, and are they convergent with other host systems? Do species within the gut microbiome vary in their temporal dynamics, and can this give us clues into ecolological differences? How does the host’s gut transcriptional landscape change in concert with the microbiome? And, is the microbiome disturbance that is widely observed in wild bumblebee populations due to individual senescence? To address these questions, we conducted a cross-sectional microbiome and transcriptomic survey of *Bombus impatiens*, focusing on dynamics during the adult stage of workers. We used high-temporal-resolution sampling and a variety of molecular methods (16S rRNA amplicon sequencing, metagenomics, qPCR, RNAseq) to provide a detailed characterization of microbiome succession and gut processes over the lifespan. Our findings develop bumblebees as a case study with which to compare dynamics with other social insects and hosts generally, and have implications for microbiome disturbance and bumblebee health.

## Results

Although our study focuses on the adult stage, we also sampled *B. impatiens* larvae to evaluate whether symbionts may persist through metamorphosis to colonize the adult gut (5). Larval microbiomes are dominated by *Lactobacillus* and *Apilactobacillus* (Fig. S7), species also present in the gut of adult worker bees (Fig. 2A). Despite this overlap, microbiomes are largely restructured across metamorphosis, with other adult-associated bacteria nearly absent in larvae (mean proportions: *Schmidhempelia*, 9.79 x 10^-4^; *Gilliamella*, 4.52 x 10^-5^, *Snodgrassella*, 3.99 x 10^-3^). Newly emerged adults (< 24 hours post-emergence) have very few bacteria in either the midgut or hindgut (Fig. 1A). 16S amplicon profiles show large proportions of reagent contaminants, such as *Burkholderia*, the most abundant taxon in our extraction blanks (Fig. S2), further indicating a scarcity of bacteria in these bees’ guts (68). As the adult bees mature, the gut bacterial community exhibits logistic growth, stabilizing after approx. 4 days, with much higher abundances in the hindgut than in the midgut (Fig. 1A). Therefore, we focus on the adult worker hindgut in the following analyses, which involve commercially reared colonies unless otherwise noted. Alpha diversity also increases quickly in young bees, from a monodominance of *Schmidhempelia* to a stable community of ∼8 bacterial groups (Fig. 2A). There was no evidence of a change in absolute abundance or alpha diversity in old bees (Figs. 1A, 1B). These patterns are highly consistent among the three replicate colonies (Figs. 1, 2). Community composition also changes with age (F= 17.7, p < 0.001) (Fig. 1C), and only weakly differs between the three replicate colonies (F = 2.18, p = 0.026).

**Figure 1.**
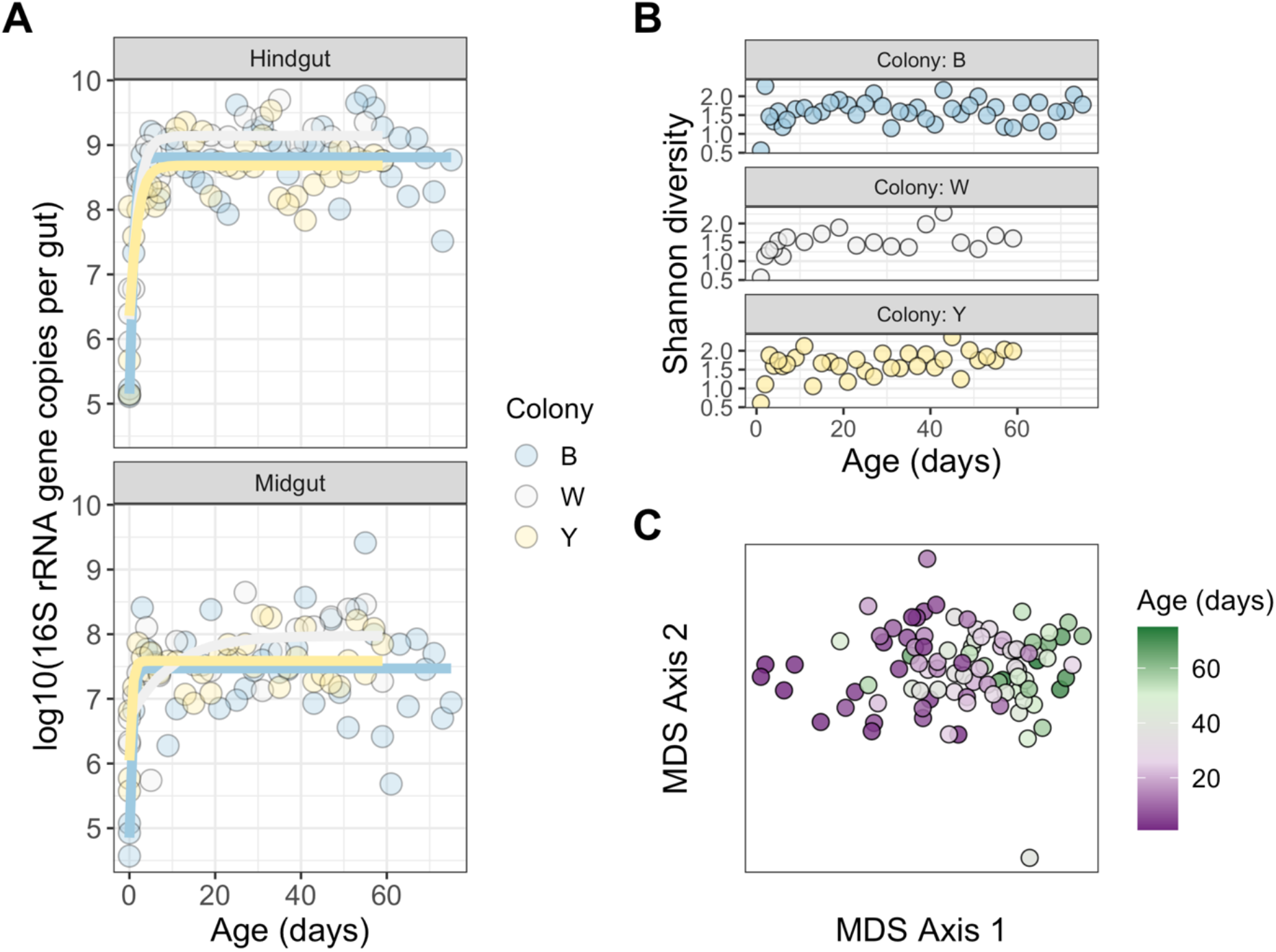
Changes in gut microbiome abundance, diversity, and composition over the worker lifespan. A total of 103 bees were sampled, consisting of 44, 23, and 36 from colonies B, W, and Y, respectively. A) qPCR-based measurements of bacterial titer as a function of age, showing patterns for each replicate colony and gut region. Solid lines are logistic curves fitted to the data. B) Alpha diversity of bacterial communities in hindguts of ≥1-day-old bees only, characterized by 16S rRNA gene sequencing. C) Beta diversity of the same hindgut samples visualized as an ordination (non-metric multidimensional scaling) of Bray-Curtis dissimilarities.

**Figure 2.**
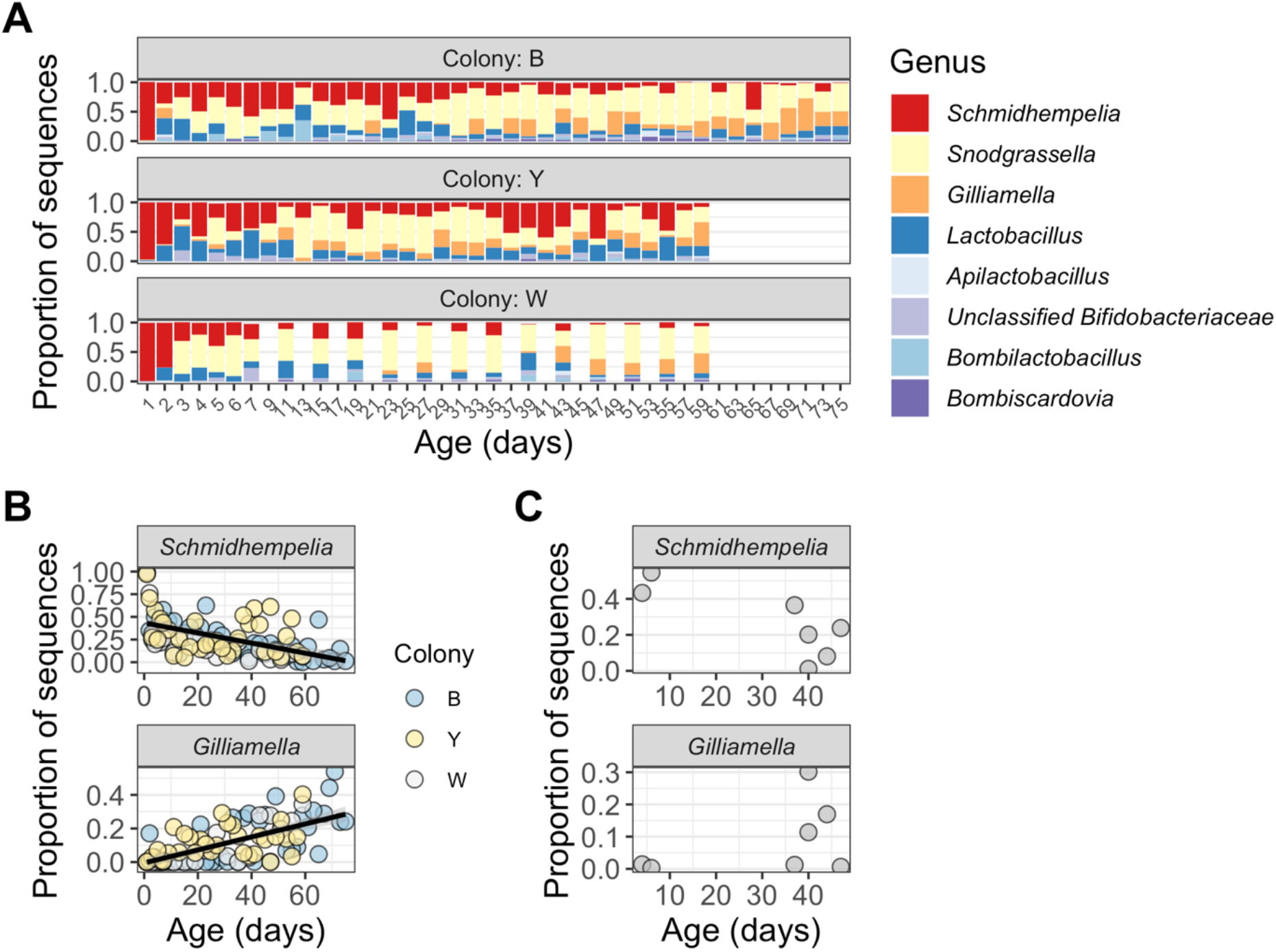
Dynamics of dominant gut microbiome taxa over the lifespan. A) 16S-based relative abundances of the top genera in hindgut samples. A total of 94 bees are shown, consisting of 41, 20, and 33 from colonies B, W, and Y, respectively. One taxon belonging to the Bifidobacteriaceae was not classified to the genus level using the SILVA database. Also note that the sampling interval varied among the three colonies (see Methods and Fig. S1). B) 16S-based relative abundances (in the same hindgut samples) for *Schmidhempelia* and *Gilliamella*, the only two taxa that varied significantly with age. Lines are linear models fitted to the data, with 95% confidence intervals in gray. C) Relative abundances of *Schmidhempelia* and *Gilliamella* in whole guts of seven workers from three *Bombus impatiens* colonies reared from wild queens.

Despite exposure to microbes present in the diet and rearing environment, gut microbiomes of workers from both commercial *B. impatiens* and wild-queen-derived *B. impatiens* and *B. ternarius* colonies are almost entirely dominated by the core, host-specialized bacterial taxa known to be prevalent in bumblebees (35) (Figs. 2A, S3). After the colonization phase, hindgut microbiome composition is generally stable throughout the adult stage (Fig. 2A), with only two taxa changing in relative abundance: *Schmidhempelia* steadily decreases with age (t = -4.48, p < 0.001) while *Gilliamella* increases (t = 7.06, p < 0.001) (Fig. 2B). As total community abundance is stable over this period (Fig. 1A), these changes reflect actual decreases and increases in population sizes. We observe the same pattern in *B. impatiens* colonies reared from wild queens (Fig. 2C). Microbes previously observed in microbiome-disrupted bumblebees, such as Enterobacteriaceae (54, 69, 70), *Fructobacillus* (55, 71), and fungi (72), are virtually absent in the 16S amplicon datasets, including commercial bee midguts (Fig. S8) and hindguts (Fig. 2A) and wild-queen-derived colonies (Fig. S3). The single exception is a male bee from one of the latter colonies, which has a large proportion of *Klebsiella* (Enterobacteriaceae) and fungal sequences (Fig. S3).

Metagenomic data provide further support for a *Schmidhempelia*/*Gilliamella* transition; using read mapping to metagenome-assembled-genomes (MAGs) as a measure of bacterial abundance, we find the same switch with age (*Schmidhempelia*: t = -3.27, p = 0.002; *Gilliamella*: t = 4.62, p < 0.001) (Fig. 3). Metagenomes also show that gut microbiomes are dominated by core bacteria. All of the MAGs belong to bee-specific bacterial taxa (Table S1); SSU rRNA genes from fungi (homologous to bacterial 16S rRNA genes) are generally rare relative to those from bacteria, though with elevated proportions in a few of the youngest and oldest bees in our sample set (Fig. S6). SSU rRNA genes from other non-bacterial microbes are practically non-existent. We also detect diet-derived plant sequences. Excepting the youngest bees, proportions of plant sequences are generally low and do not show any clear trends with age (Fig. S6).

**Figure 3.**
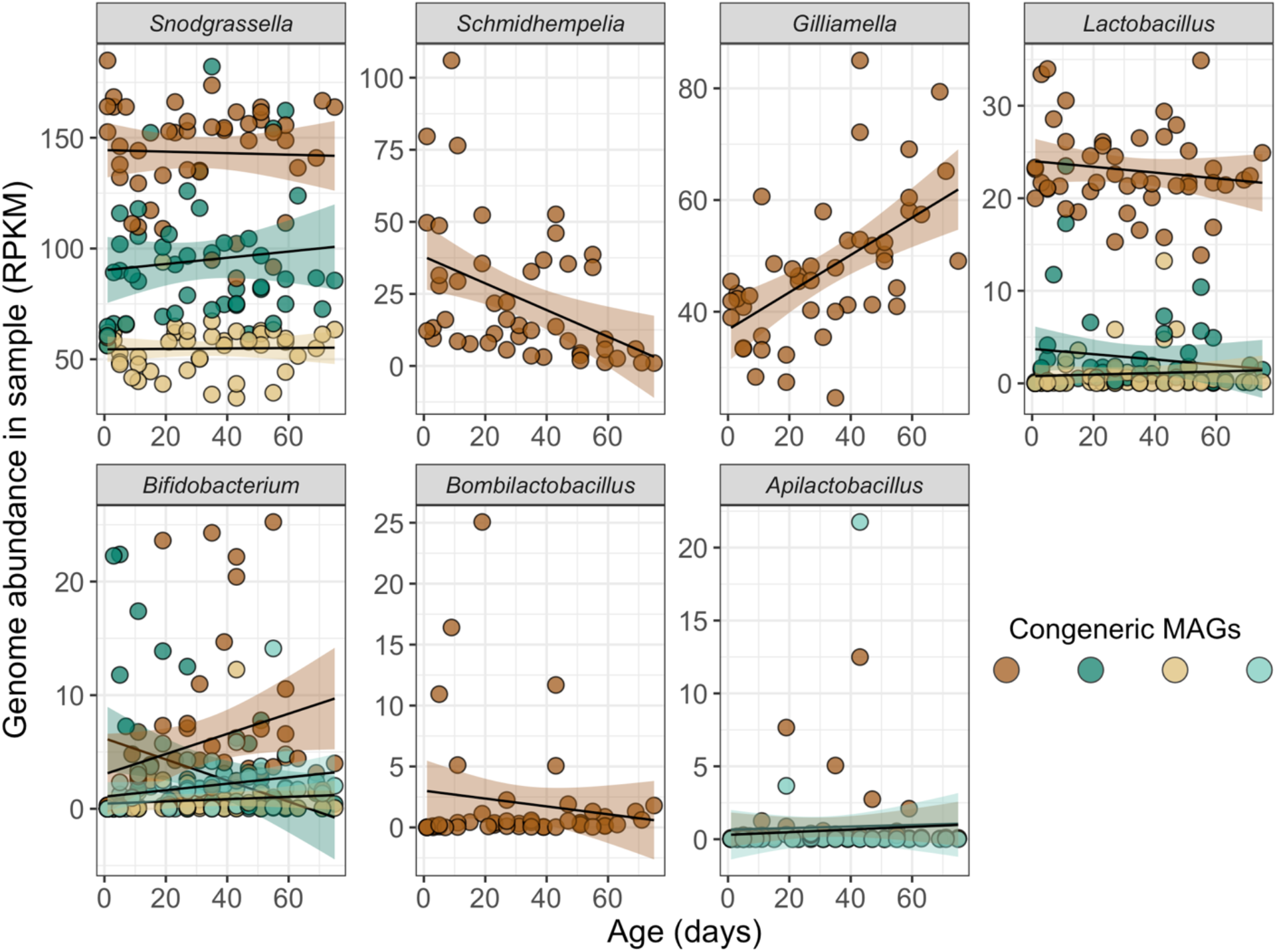
Coverage-based abundance estimates of all metagenome-assembled genomes (MAGs) in 46 worker hindgut samples from the three commercial colonies. Abundance for a given sample is normalized to sequencing depth and MAG size, by measuring reads per kilobase per million mapped reads (RPKM). Lines are linear models fitted to the data, with 95% confidence intervals. Some genera contained multiple MAGs with < 98% average nucleotide identity; in these cases, congeneric MAGs are shown in different colors. MAGs are listed and described in Table S1.

Analysis of amplicon sequence variants (ASVs), the finest level of resolution available with our 16S sequencing approach, shows that the major core taxa comprise only a single ASV generally ubiquitous across samples (Fig. S4). We used metagenomic data to reveal further layers of diversity beyond ASVs. Some (but not all) of the major bacterial groups comprise multiple MAGs with < 98% ANI [“subspecies”, following (73)] (Fig. 3, Table S1). Using inStrain, which compares single-nucleotide variants between samples’ reads aligned to a common reference (73), we find that MAGs contain additional strain-level diversity. For most MAGs, this diversity is clearly partitioned by colony, but not by age (Fig. 4, Fig. S5). All of the MAGs from Gram-negative bacterial taxa (*Snodgrassella, Schmidhempelia, Gilliamella*), but only some of those from Gram-positive taxa, are more likely to be shared within than between colonies (FDR-adjusted p < 0.05) (Fig. S5). We also used metagenomic data to examine *in situ* population-average replication rates, focusing on the two taxa that shift with age. *Schmidhempelia* has much lower replication indices in colony B (posthoc pairwise contrasts: W vs. B, t = 10.69, p < 0.001; Y vs. B, t = 14.29, p < 0.001; W vs. Y, t = 1.35, p = 0.38) (Fig. 5). There is also a weak negative effect of age on *Schmidhempelia* replication (t = -2.65, p = 0.013). *Gilliamella* replication indices do not significantly differ between colonies (t = -1.32, p = 0.22) or due to age (t = 0.197, p = 0.89) (Fig. 5), although sample sizes are also smaller due to lower coverage.

**Figure 4.**
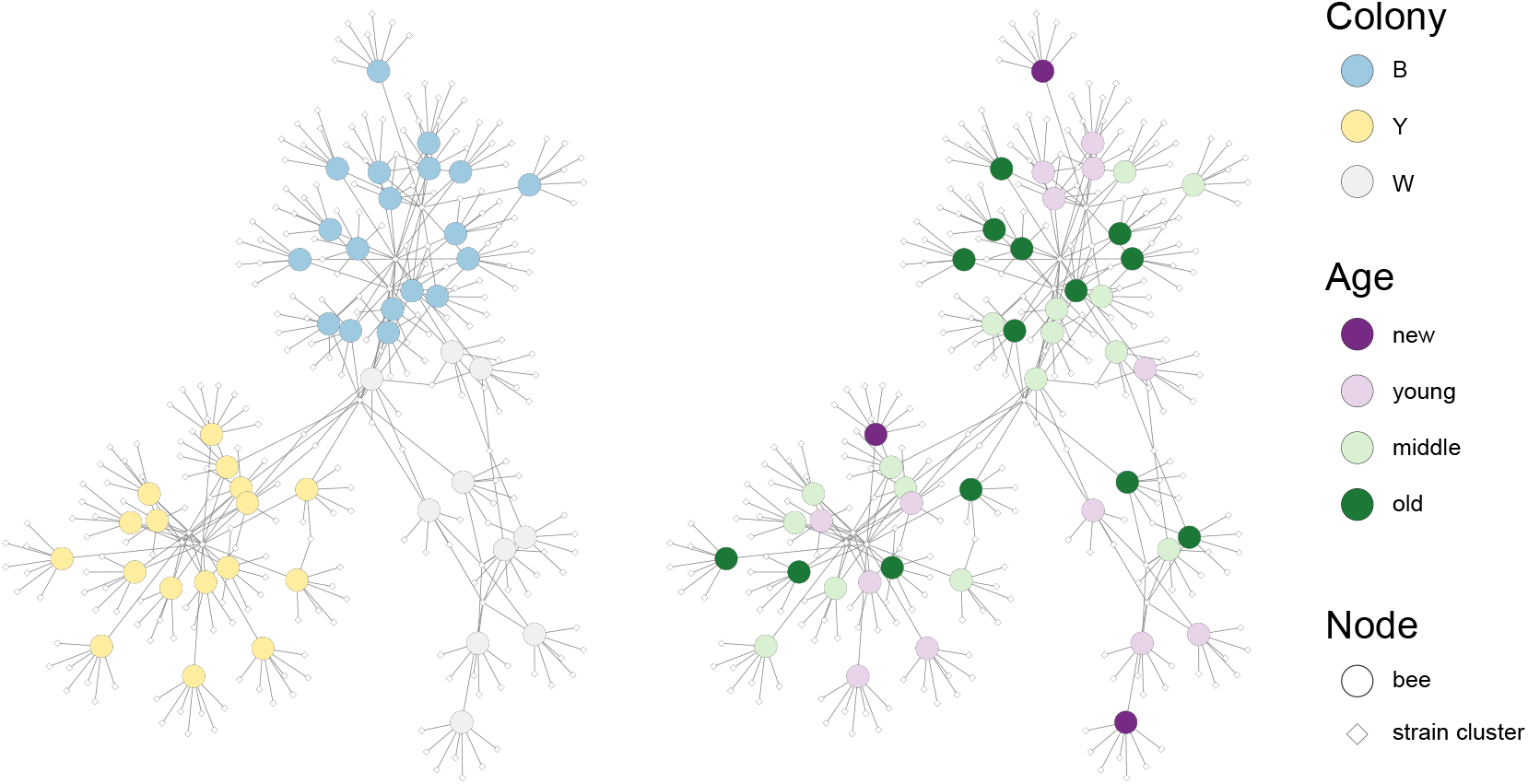
Networks of bacterial strain composition in the 46 worker hindgut metagenomes, showing sample grouping by colony versus age class. Strain clusters from all MAGs are shown; strain sharing within and between colonies is shown for each MAG individually in Fig. S5. Clusters are derived from hierarchical clustering of pairwise comparisons of population ANI, a metric calculated by inStrain (see Methods).

**Figure 5.**
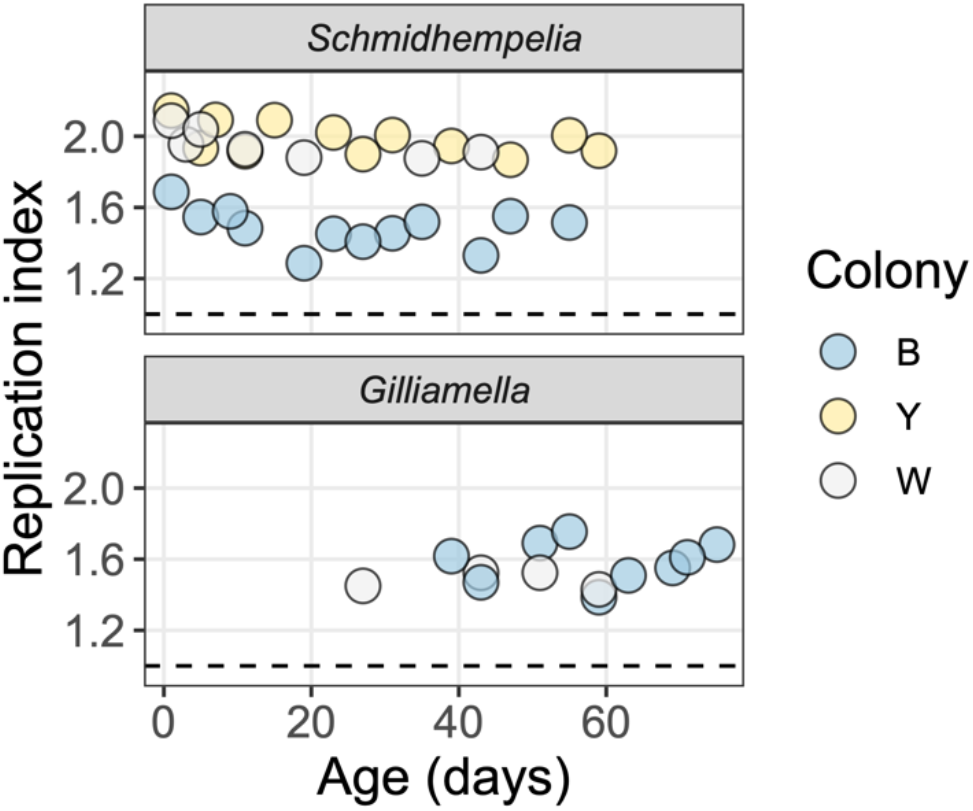
Instantaneous population-average replication rates, estimated for *Schmidhempelia* and *Gilliamella*, the two taxa that vary in abundance with age. A replication index value of 1.5 corresponds to half of the cells making one copy of their genome; with a value of 2, all cells are making one copy [see ref. (131)]. However, note that these are population averages, and bacteria can make multiple copies of their genome simultaneously. Some data points are missing due to low coverage of the MAG in a given sample.

Host gene expression profiles in the hindgut change as bees mature and reach “middle age” (∼3-6 weeks old) (Fig. 6A). Between newly emerged and young bees, and young and middle-aged bees, there are 2696 and 6136 differentially expressed genes (DEGs), respectively. Thereafter, gut gene expression profiles do not change with age in a consistent way (Fig. 6A): there are zero DEGs comparing middle-aged and old bees. Of the immunity-related genes we analyzed in more detail, most show low levels of expression in newly emerged bees, with upregulation in older age cohorts (Fig. 6B). Three of the four antimicrobial peptide genes, as well as dual oxidase [which generates reactive oxygen species (ROS) (74, 75)], increase in expression as bees mature. Catalase, which degrades ROS to maintain redox balance (76), is highly expressed in newly emerged bees—possibly to prevent self-harm in the absence of abundant microbial cells (60)—and is subsequently downregulated. Overall, immune gene expression either increases or remains stable with age (Fig. 6B).

**Figure 6.**
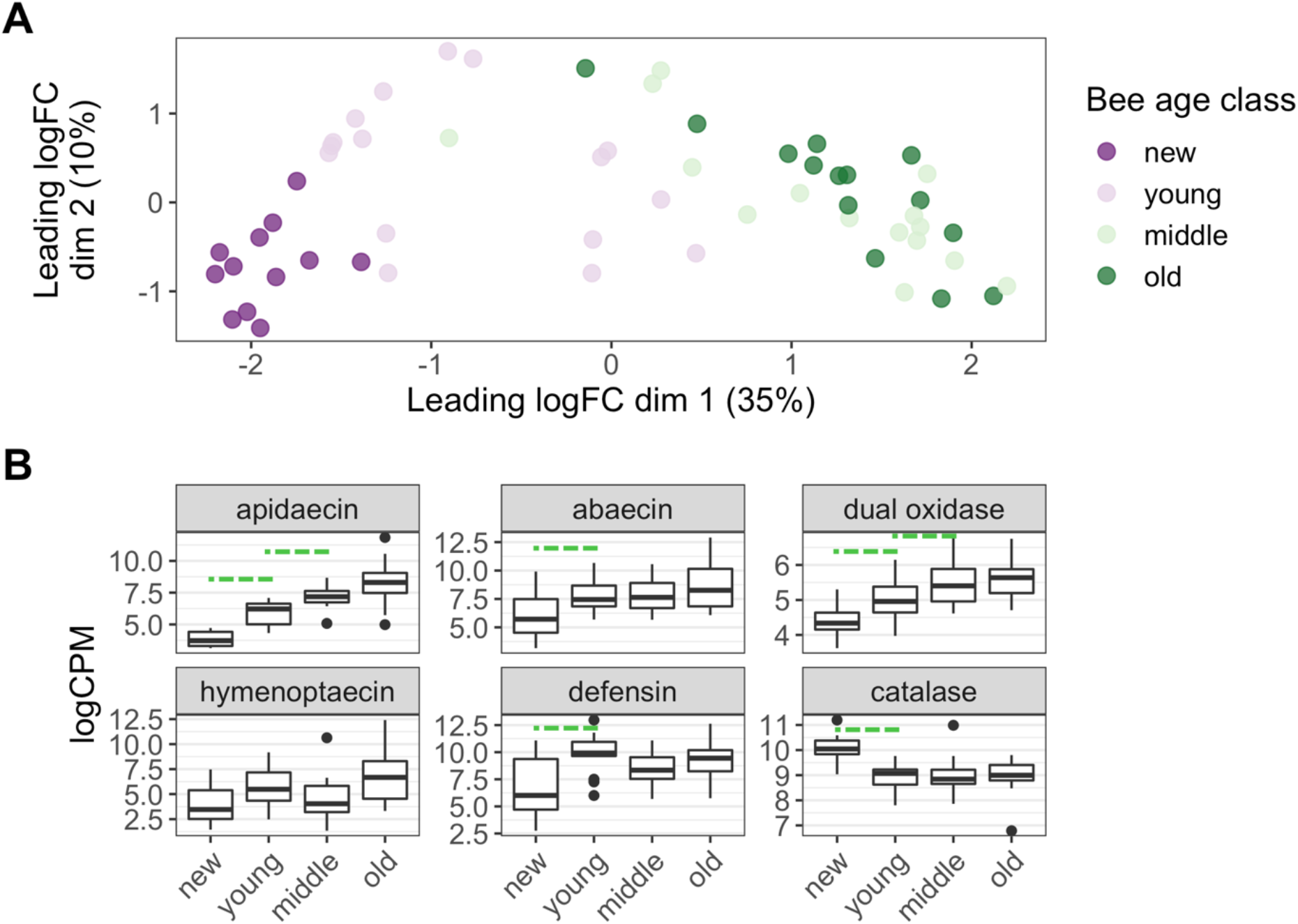
Dynamics and stability of host hindgut gene expression over the lifespan. Ages and sample sizes of age classes: new: 0-1 days, N = 12; young: 3-19 days, N = 16; middle: 23-43 days, N = 15; old: 47-75 days, N = 14. A) Principal coordinates analysis showing similarity in gene expression profiles between bees of different age classes. Similarity is quantified as leading log2-fold changes, which are defined as the quadratic mean of the largest log2-fold changes between a pair of samples. Boxplots display differences along the first axis between newly emerged, young, and middle-aged bees; middle-aged and old bees overlap, with no genes differentially expressed between them. B) Expression levels of key immunity genes normalized to library size (log2 counts per million) over bee age. Dashed lines show significant differences in expression between age classes (FDR-adjusted p < 0.05).

## Discussion

### Assembly

In adult bumblebees, microbiome assembly appears to be an example of primary succession. Amounts of bacterial DNA in newly emerged adult guts are very low (Fig. 1A), and previous work finding these guts generally devoid of culturable bacteria (77, 78) suggests that this DNA may derive from nonviable cells. Larvae harbor *Lactobacillus* and *Apilactobacillus* (Fig. S7), taxa also present in adult guts (Fig. 2A). Although transmission through metamorphosis is theoretically possible, these bacteria may instead be cleared during pupation and reacquired from the nest environment by newly emerged adults (5). Other dominant bacteria in adult guts such as *Schmidhempelia, Snodgrassella*, and *Gilliamella* were largely or completely absent from larvae (Fig. S7), indicating *de novo* colonization of adults.

Coupled with a low expression of immunity genes (Fig. 6B), the low abundance of pathogen-protective (79, 80) core gut bacteria suggests that newly emerged adults are particularly vulnerable to microbiome disruption. Similarly, human infants are prone to infections while their immune system, microbiome, and gut microenvironment mature (81). The microbiome disruption phenomenon widely observed in field-collected bumblebee workers as well as queens (35) may begin during the assembly phase. In *Bombus griseocollis*, workers do not leave the nest for the first couple of days after emergence; most activities, including foraging, begin by the fourth or fifth day (52). By this point, the core gut microbiome is established and immunity has increased (Fig. 1, Fig. 2, Fig. 6B). The timing of the onset of foraging therefore limits direct exposure to stressors during this vulnerable period. However, environmental microbes and chemicals are present in food stores and other substrates, presenting an opportunity for microbiome perturbations even in bees restricted to the nest.

Microbiome assembly dynamics in bumblebees are both predictable and convergent with other hosts. Temporal patterns of microbiome abundance, diversity, and composition (Fig. 1) are highly similar among replicate colonies. Moreover, these patterns are evident despite our cross-sectional study design, suggesting that temporal variation in microbiomes outweighs interindividual variation. Successional patterns showed similarities with those observed in honeybees (22) and human infants (4, 20, 82, 83) and, more generally, to heterotrophic microbial communities supplied with external carbon sources (3). However, there are also marked differences with gut microbiome assembly in other invertebrates, such as flies (*D. melanogaster*) and nematodes (*C. elegans*). In these hosts, bacterial colonization is highly stochastic and can lead to microbiome compositions that are stably distinct among individuals (13, 14, 19). These hosts also generally harbor non-host-restricted, flexible, environmentally acquired gut microbiomes (84). In contrast, the symbiosis between social bees and their gut microbes is ancient and specific (34, 41). By living in dense colonies, social bees enrich their local environment with core symbionts, favoring predictable assembly. Functional redundancy among bacterial species may also be lower in social bees than flies and nematodes, possibly selecting for stronger host control over microbiome establishment.

Despite predictable assembly at the community level (Figs. 1, 2), we also observe evidence for stochastic colonization at the strain level. Strain-level diversity is clearly partitioned between the three replicate colonies (Fig. 4), a pattern not evident in the ASV (Fig. S4) or subspecies (Fig. 3) data. Notably, all Gram-negative bacterial genomes exhibited significant colony partitioning, while only some of the Gram-positive genomes did so (Fig. S5). Gram-positive bacteria may be more likely to survive outside the host, facilitating dispersal among colonies. Similarly, Gram-positive gut bacteria of honeybees can be transmitted via hive surfaces, with less reliance on social contact than Gram-negative species (22). Differences in social structuring among mammalian microbiome members have also been linked to bacterial physiology (85, 86).

There are multiple potential explanations for the origin of the colony-partitioning pattern. One is an interaction between host and symbiont genotypes (77). There may also be genotype-by-environment effects; to give one example, bee colonies of different sizes may have different thermoregulatory capacities and temperatures (87); this could act as an ecological filter for strains with different thermal tolerances (78). In addition to intrinsic physiological differences between strains, differences in temperature or other environmental factors may explain why the inferred replication rates of *Schmidhempelia* differed substantially between colonies (Fig. 5). A final explanation is founder (or foundress) effects. Bumblebee colonies are initiated by a single foundress queen, who is the source of gut symbionts for her offspring (35, 56). A diverse pool of strains may be stochastically sorted into a single foundress queen’s gut, with the established population resistant to subsequent invasion [i.e., priority effects (2)]. This process may be analogous to the neutral bottlenecking described for bacterial strain partioning among skin pores (88) or the stochastic colonization of individual guts of flies and nematodes (13, 14, 19).

### Maintenance

While microbiome abundance and composition generally stabilize after the colonization phase, the ratio of two of the core symbiont species, both members of the family Orbaceae, continues to shift. *Schmidhempelia* progressively declines in relative abundance, while *Gilliamella* increases. This pattern is evident in both the amplicon (Fig. 2B), and metagenome datasets (Fig. 3), and across the three replicate colonies. We observe the same pattern in wild-derived colonies (Fig. 2C), suggesting that it is a common feature of microbiome succession in *B. impatiens*. Such age-based shifts could be broadly conserved among social bees, as a similar pattern occurs in honeybees for *Gilliamella* and *Frischella*, another Orbaceae species (22). Gene expression profiles in the hindgut are also dynamic up to ∼3-6 weeks of age (“middle age”), with many differentially expressed genes between newly emerged, young, and middle age (Fig. 6A). Multiple genes involved in production of antimicrobial peptides (AMPs) and reactive oxygen species, key components of gut epithelial immunity (74, 75), increase in expression over this time frame (Fig. 6B). These microbiome and immunity dynamics appear to be intrinsic properties of aging in *B. impatiens* workers, as they occur despite continuous food availability and static environmental conditions in the laboratory. As hypothesized for temporal patterns of systemic immunity in bumblebees, they may represent a plastic adjustment of host defense, evolved in response to increasing infection risk with age (67). An alternative hypothesis is that increasing immune gene expression represents increasing inflammation with age, a common feature of animal immunosenescence (61). In *D. melanogaster*, increased AMP expression with age is linked to increased gut bacterial load and to deteriorating gut integrity (8, 29). However, total gut bacterial load in bumblebees is stable (Fig. 1A), and the only taxon that increases in abundance is *Gilliamella* (Fig. 2B, Fig. 3), one of the core bumblebee-specialized symbionts (35). While *Gilliamella* may induce bee AMP expression (60), such a response with age could be interpreted as a sign of strengthening, as opposed to deteriorating, immunity.

The temporal dynamics of *Schmidhempelia* and *Gilliamella* point to distinct life history strategies, perhaps exemplifying the competition-colonization trade-off shown in various microbial communities [e.g., (89–91)]. For example, *Schmidhempelia* may be a pioneer colonizer or ruderal (42), one that is good at dispersing to and exploiting unoccupied gut habitat. *Gilliamella* may be a better competitor, successfully excluding *Schmidhempelia* with time. The nature of this competition remains to be determined. *Schmidhempelia* replication rates are generally stable with age after the colonization phase (Fig. 5), suggesting that declines in population size are driven by increased mortality over time, rather than by a dwindling resource supply slowing replication. Increased mortality could be due to interference competition, where *Gilliamella* directly antagonizes *Schmidhempelia* using type VI secretion systems [possessed by both species (53, 92)] or other means. It could also be due to apparent competition, where *Gilliamella* growth induces increased expression of host immune responses (Fig. 6B) that are more harmful to *Schmidhempelia* than to *Gilliamella* (60). Although the mechanisms are unknown, our data at least support the existence of variation in life history strategies within the gut microbiome. Such differences are poorly understood, but are likely to be important drivers of coexistence and community function.

Both microbiome and transcriptome dynamics slow as bees enter old age, contrasting with dynamics observed between life stages (Fig. S7) and observed earlier in the adult stage (Figs. 1, 6). Total microbiome abundance is stable in old bees (Fig. 1A), and there is no evidence of microbiome disruption—with the exception of a single male from a wild-derived colony (Fig. S3)—or loss of any symbionts besides *Schmidhempelia* (Fig. 2). All bees were reared indoors, and indoor-reared bumblebees have been shown to have lower gut microbiome diversity (35, 69, 70). However, the bees studied here were exposed to non-core microbes in their food and rearing environment, and previous work has documented occasionally large numbers of Enterobacteriaceae and other non-core bacteria in indoor-reared *B. impatiens* when exposed to stressors (93–95). The microbiome stability we observe in old bees indicates a lack of intrinsic senescence processes that would disrupt core symbionts and allow invasion, rather than simply a lack of exposure to non-core microbes. Bumblebees therefore contrast with humans (23–26), as well as other animals such as flies, mice, and fish (17, 27–29), which exhibit microbiome senescence (or at least community-wide shifts during aging) even when reared in the laboratory. Our data also weigh against the hypothesis that individual senescence underlies the microbiome disturbance observed in wild bumblebee populations. As mentioned above, it is the youngest bees who appear to be the most vulnerable. These results support previous work finding microbiome disruption to be concentrated in young bumblebees (70).

There are many potential proximate causes of microbiome stability in old age. Communal living may buffer microbiome disturbances by providing a continuous source of microbes that can be socially transmitted (85, 86, 96, 97). In our experiment, diet was kept constant, and bees appear to consume pollen even in old age based on the presence of plant DNA in metagenomes (Fig. S6) and observations of gut color. Our transcriptomic data also suggest stable physiological and immunological processes in the gut, as no genes were consistently up-or down-regulated between middle-aged and old bees (Fig. 6A). In addition to inoculation from nestmates, a steady resource supply, structural integrity, and elevated immune responses in the gut likely help maintain stable core microbiomes. One caveat is that these bees were not able to fly, a factor that should be addressed in future work. Bee flight is metabolically costly, reduces lifespan, and affects systemic immune responses (98–100). Free-foraging honeybee workers do exhibit changes in gut microbial abundance, composition, and replication rates with age (15, 22, 45–47). On the other hand, as noted earlier, it is difficult to differentiate the contributions of aging versus changes in task performance in honeybees.

Overall, bumblebee gut microbiome senescence and immunosenescence appear to be either absent, or compressed into such a short window that we did not observe it. We suggest that this may be explained by the unique selection pressures that accompany eusociality. Evolutionary theories of aging suggest that in a non-social host organism, i) selection against late-acting, deleterious variants—either host alleles or microbes—should be weak, and ii) such variants may trade off with early-life, pre-reproductive benefits (101–103). The situation is different in bumblebee workers, which are (usually) nonreproductive and which often complete their entire life cycle before colony reproduction (64). In these workers, maintenance of microbiomes and immunity should be under strong selection even in old age, given their expected effects on inclusive fitness, that is, overall colony reproductive success. Core gut symbionts may contribute indirectly (e.g., via nutrition) to worker performance—brood care, foraging, defense, etc.—which in turn will affect production of new queens and males at the end of the colony cycle. Workers may also benefit their reproductive siblings (the new queens and males) by acting as a vector for core symbionts, and not for pathogens or parasites. Microbiomes of at least some other highly social animals do not appear to become destabilized in old age (32, 33), raising the question of whether group living contributes to differences in microbiome senescence. In general, organisms display diverse patterns of mortality and reproduction with age (104). We should expect such diversity to extend to microbiome dynamics, and seek to understand what drives it.

## Conclusions

Even in the relatively simple gut microbial communities of laboratory-reared worker bees, we see a complex assortment of temporal patterns that differ between symbiont taxa, vary with phylogenetic scale, and decelerate as hosts age. Some of these patterns are convergent with those in other hosts. At the community level, assembly is predictable and similar to the dynamics of human infant gut microbiomes; at the strain level, assembly appears similar to the stochastic colonization dynamics observed in flies and nematodes. We also find unique temporal patterns that contrast with those in other hosts: in bumblebee workers, neither gut microbiomes nor gut immunity appear to senesce. This stability may be due to the important contributions of each to inclusive fitness, even in old age. Temporal dynamics differ markedly among bacterial symbiont species, suggesting distinct ecological strategies within the microbiome for colonization and persistence. Many of the patterns we observe would be undetectable by 16S rRNA gene sequencing, emphasizing the need to use quantitative and higher-resolution methods to study microbiome dynamics. We also characterize the transcriptomic landscape of the bumblebee gut, finding that expression of genes involved in immunity (and other processes) changes in similar ways to the microbiome over host age—likely due to bidirectional feedbacks or to common selection pressures acting on both. A priority for future work is to determine the mechanisms underlying these microbial and immunological dynamics, and assess functional consequences for bumblebee health and pollination services.

## Materials and Methods

### 1. Bumble bee rearing

For the main study, three commercially reared bumble bee (*Bombus impatiens*) colonies were obtained from Koppert Biological Systems and reared in the laboratory. Upon arrival, all of the cocoons (containing worker pupae) present in each colony were moved to separate containers in a 35 °C incubator. We monitored the cocoons daily, and marked all newly emerged adult worker bees with numbered tags, affixed with wood glue to the thorax. Tagged bees were then returned to their colony of origin. Three newly emerged bees per colony were sampled (see below) instead of returned to the colony. To maintain colonies, we provided non-sterile pollen dough (ground pollen mixed with syrup) every 3-4 days. Non-sterile sucrose syrup (50% w/v) was provided ad libitum through an enclosed foraging area connected to the main nest.

We used cross-sectional sampling to measure changes in gut microbiomes and transcriptomes over the worker lifespan (Fig. S1). For the first week of adult life, we sampled one bee per colony per day, in order to have higher temporal resolution for the colonization phase, which we expected to be dynamic. Thereafter, for colonies B and Y—which had more tagged bees available, because more pupae were present—we sampled one bee every other day in age (e.g. 9, 11, 13 days old). For colony W, sampling occurred every fourth day in age. Sampling entailed anesthetizing bees on ice and removing the gut with 70% ethanol-sterilized forceps. The midgut and hindgut were separated at the pylorus and each stored in 0.1 ml DNA/RNA Shield (Zymo) at -80 °C until nucleic acid extractions.

Sampling continued until all of the originally tagged bees had either died or been collected—up to 59 days old (colonies Y and W) or 75 days old (colony B) (Fig. S1). These maximum ages are similar to, or greater than, the average lifespan for indoor-reared workers of *Bombus impatiens* (105, 106) and other *Bombus* species (107, 108). They greatly exceed the average lifespan of free-foraging bumble bee workers (109–111).

A smaller set of samples were collected from colonies reared from field-collected queens of *B. impatiens* (3 colonies) and *B. ternarius* (1 colony). Queens were collected from New Hampshire, USA (*B. impatiens*: 44.221788, -71.735138; *B. ternarius*: 44.221034, -71.774747). They were then reared in small Ziploc containers in the closet of a private residence at ∼60% relative humidity and at 28 °C. The colonies were fed pollen and nectar as described above. Newly emerged bees were tagged and returned to the colony, and combined midgut and hindgut samples were collected from younger (4-14 days old) and older (37-47 days old) workers and males. Samples were stored in 95% ethanol at -20 °C. Finally, we also sampled 11 larvae from two additional commercial *B. impatiens* colonies. Larvae were stored in 95% ethanol at -20 °C.

### 2. Nucleic acid extractions and qPCR

All samples were homogenized with a sterile pestle prior to extractions. For hindguts sampled from the commercial colonies, we extracted both DNA and RNA using the Zymo Quick-DNA/RNA kit, following the manufacturer’s protocol. For all other samples we extracted only DNA using the ZymoBIOMICS DNA kit. Six extraction blanks and three cross-contamination controls (0.l ml of a OD600 10.0 suspension of *Sodalis praecaptivus* cells in PBS) were included alongside the gut samples.

Bacterial titers were measured by SYBR Green-based quantitative PCR targeting the 16S rRNA gene (with universal 27F/355R primers), as described in ref. (22). Absolute copy numbers were calculated using standard curves generated from serially diluted plasmid DNA carrying the target gene. Estimates of copy numbers per gut sample were calculated by multiplying values from qPCR reactions (containing 1 μl template) by the volume of gDNA eluted from each extracted sample.

### 3. Library prep and sequencing

For 16S rRNA gene sequencing, gDNA (excepting larval samples) were first PCR-amplified using universal primers targeting the V4 region (515F/806R) and conditions as detailed in ref. (112). Addition of dual-indexed barcodes, magnetic bead purifications, and additional library preparation steps also followed the protocols in ref. (112). Libraries (including the extraction blanks and three PCR no-template controls) were pooled and sequenced on an Illumina iSeq with 2 x 150 chemistry. Samples were split among three separate sequencing runs, as listed in the supplementary metadata file. For larvae, library prep and 16S rRNA gene (V4 region) sequencing (Illumina NovaSeq 2 x 250) were conducted separately by Novogene.

A total of 57 hindguts from the commercial colonies, spanning the range of ages in our sample set (including newly emerged bees), were selected for RNAseq. Library prep for host mRNA sequencing was conducted by Novogene using the NEB Next Ultra II RNA library prep kit. Libraries were sequenced on an Illumina NovaSeq with 2 x 150 chemistry, resulting in an average of 23.7 M raw paired-end reads per sample. The same set of hindguts used for RNAseq were initially selected for shotgun metagenomics, excepting the newly emerged bees, which had very low amounts of bacterial DNA. Five gDNA samples did not pass QC, and three of these were replaced by other hindgut samples from bees “adjacent” in age, for a total of 46 samples. Library prep was conducted by Novogene, using the NEB Next Ultra II DNA Library prep kit. Libraries were sequenced on an Illumina NextSeq with 2 x 150 chemistry, with an average of 22.4 M raw paired-end reads per sample.

### 4. Bioinformatic analyses

16S rRNA gene amplicons from the three iSeq runs were combined for data processing. Adapters and primers were removed using cutadapt (113). Sequences were then quality-filtered, trimmed, and denoised to generate amplicon sequence variants (ASVs) by DADA2 (114). Only the forward reads were used, as the reverse reads were poor quality. Taxonomy was assigned to ASVs using the SILVA v. 138.1 database (115). Further data processing and analysis was conducted in R v. 4.1.1 and followed the general approach described in ref. (116). ASVs with < 100 total sequences across all samples were removed. ASVs classified as mitochondria or chloroplast were also removed (mean 3.6% of sequences). *Sodalis praecaptivus* was almost nonexistent in gut samples (mean 0.008% of sequences), suggesting minimal cross-contamination. One unclassified ASV (ASV_77) was highly abundant in a single bee; as its sequence was 100% identical to *Aspergillus* and *Mucor* mitochondrial 16S rRNA gene sequences in NCBI, it was reclassified as Fungi (see Fig. S3). Libraries were rarefied to 2200 sequences, leaving out three samples with <1000 sequences (one midgut and two hindguts from newly emerged bees). Sequences from larvae were processed separately. For these data, removal of adapters and primers, and quality-filtering, were conducted by Novogene. Paired-end reads were also merged before dereplication and denoising to generate ASVs. Otherwise, processing followed the steps described above. ASVs classified as mitochondria or chloroplast comprised a mean 1.3% of sequences in larvae, and these samples were rarefied to 66258 sequences.

For the RNAseq data, we used scythe (https://github.com/vsbuffalo/scythe) to remove adapters, and then sickle (https://github.com/najoshi/sickle) was used to trim reads with a minimum quality score of 33. Reads were aligned to the *Bombus impatiens* reference genome BIMP_2.2 (117) using STAR (118). Analysis of count data in R followed the general approach of ref. (119), using limma (120) and edgeR (121) packages. Genes were filtered using the filterByExpr function, with normalization factors calculated by the TMM method. To use pairwise differential expression analyses, we grouped bees into four age classes with roughly similar sample sizes. We calculated the number of differentially expressed genes (DEGs) between age classes using linear models of log counts-per-million (log-CPM) values in limma. The design matrix (∼ 0 + age + colony), and contrasts were designed for pairwise comparisons of adjacent age classes (e.g., young versus middle-aged). DEGs were defined as genes with a p value <0.05 after false discovery rate (FDR) adjustment for multiple comparisons. To analyze expression patterns of genes that might be linked to microbiome dynamics, we focused on antimicrobial peptides and dual oxidase, which generates reactive oxygen species. Both are major components of gut epithelial immunity in insects, and are known to regulate gut microbes in other species (74, 75).

For the shotgun metagenomic data, we used cutadapt (113) to remove adapters and quality-filter reads. These reads were mapped to the *B. impatiens* reference genome BIMP_2.2 (117) using bowtie2 (122), with unmapped reads used for further processing. phyloFlash (123) was used to analyze the non-bacterial taxonomic composition of SSU rRNA genes. To assemble the data, we used megahit (124) for single-sample assemblies (125). Assemblies were binned using both MaxBin 2.0 (126) and MetaBAT 2 (127). Then, we used dRep (128) to obtain a single, dereplicated set of bacterial metagenome-assembled genomes (MAGs) with an average nucleotide identity (ANI) threshold of 98%. Three MAGs with >2% contamination and one MAG with <90% completeness were discarded, leaving a set of 15 high-quality MAGs (Table S1). These MAGs were classified using GTDB-Tk (129). For further analyses, we mapped each sample’s reads against the concatenated set of MAGs using bowtie2. The relative abundance of each MAG in each sample, normalized by sequencing depth, was measured as the number of reads per kilobase per million mapped reads (RPKM), calculated using the pileup function from BBMap (sourceforge.net/projects/bbmap/). inStrain (73) was used to resolve strain-level diversity. Specifically, we characterized strain-level clusters (generated by the inStrain compare function) and visualized their distribution across samples using cytoscape (130). We also conducted a non-clustering-based analysis of strain sharing with 99.99% population ANI as a cutoff for differentiating strains (Fig. S5). This method has a high sensitivity to detect shared strains; in calculating population ANI, single-nucleotide variants are only called when reads of both samples (mapped to a common reference) share neither major nor minor alleles at a given position (73). Finally, we used iRep (131) to estimate the instantaneous population-average replication rates for *Schmidhempelia* and *Gilliamella* MAGs in each sample.

### 5. Statistical analyses

To model changes in bacterial titer with age, we fit logistic curves to the data using the SSlogis and nls functions in R. To test whether changes in community composition were associated with age and colony, we used distance-based redundancy analysis with the Bray-Curtis dissimilarity metric, as implemented in the vegan package (132). To identify bacteria whose relative abundance changed with age after the colonization phase, we focused on only the dominant groups shown in Fig. 2, leaving aside very low-abundance taxa. Then we conducted Spearman’s correlations and adjusted p values using FDR. For the three taxa with an FDR-corrected p value < 0.05, we used linear mixed effects models to further test whether age predicted changes in relative abundance, including colony as a random effect. The latter approach was also used to test whether the relative abundance of *Schmidhempelia* and *Gilliamella* MAGs varied with age. Strain partioning by colony versus age (the four discrete age classes described above) was analyzed by the following method: for all MAGs, chi-squared tests were conducted to test whether bees belonging to the same colony or age class tended to have a higher number of shared strains; p values from these tests were corrected for multiple comparisons by FDR. To model replication indeces of *Schmidhempelia* and *Gilliamella* as a function of age and colony, we first conducted linear regressions including the interaction term; these did not provide a significantly better fit to the data than models lacking an interaction, so the results we report are from the latter. We used the glht function in the multcomp package for posthoc tests of *Schmidhempelia* replication indeces among the three colonies. R code used for statistical analyses will be made publicly available upon submission of the revised manuscript.

## Author Contributions

TJH and NAM designed the research. TJH conducted the commercial bee rearing and sampling, molecular methods, bioinformatics, and statistical analyses. AEC reared and sampled bees from the wild-queen-derived colonies. The manuscript was drafted by TJH and subsequently revised and approved for submission by all authors.

## Acknowledgments

We acknowledge Eli Powell and James Crall for technical advice and assistance, and Kim Hammond for administrative support. We also thank Liam Easton-Calabria and Sahana Simonetti for their help in constructing the home laboratory used to rear colonies from wild queens. This research was funded by a postdoctoral fellowship (2018-08156) from the USDA National Institute of Food and Agriculture to TJH, a grant from the Star-Friedman Challenge for Promising Scientific Research to AEC, and a NIH grant (R35GM131738) to NAM.

## Supplementary Table and Figures

**Table S1.**
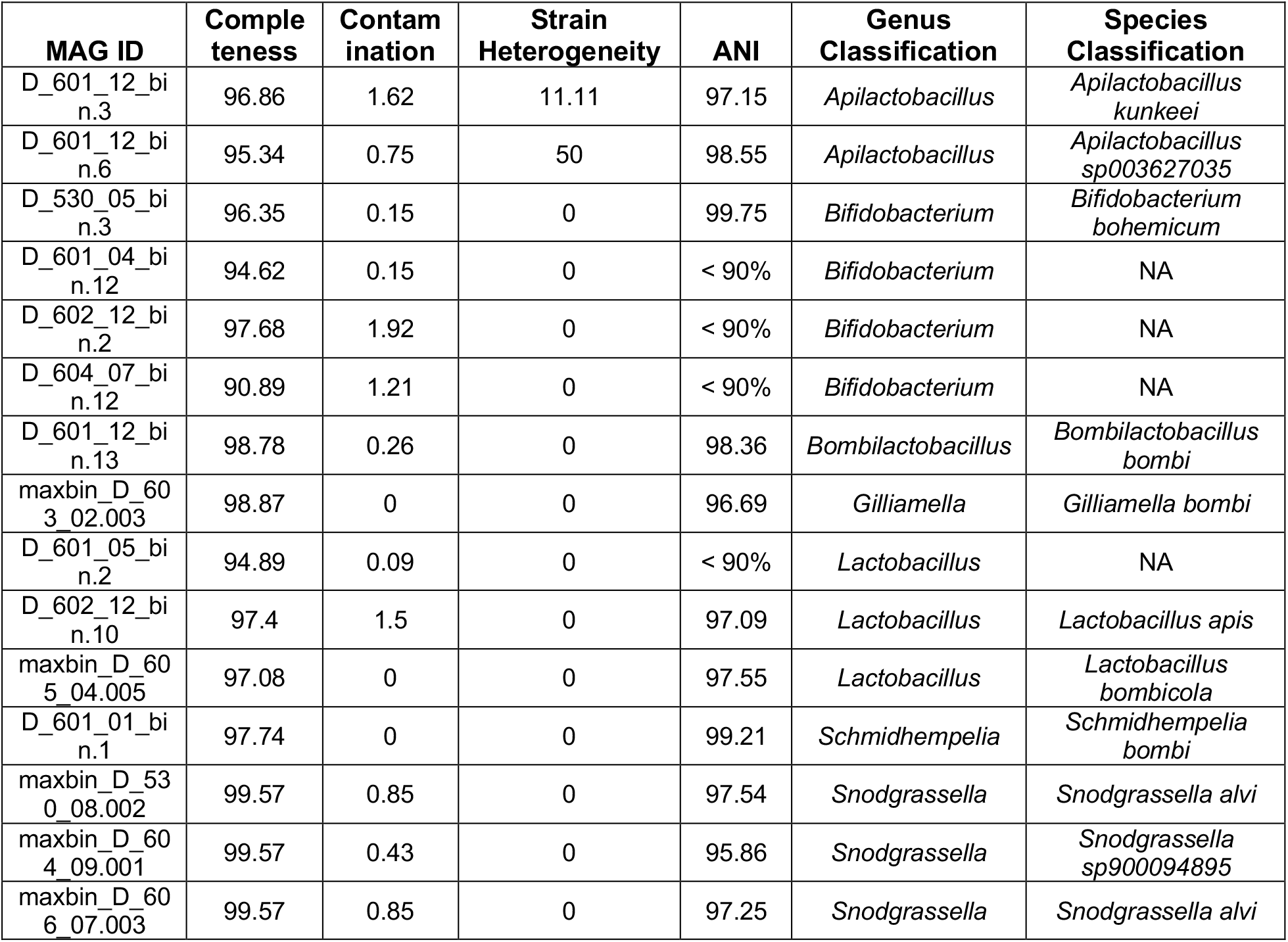
The dereplicated set of high-quality metagenome-assembled genomes (MAGs). Completeness, contamination, and strain heterogeneity are calculated by checkM using conserved single-copy genes. Taxonomic classifications are from GTDB-Tk. ANI refers to the % average nucleotide identity of the MAG to the closest reference genome in the Genome Taxonomy Database.

**Figure S1.**
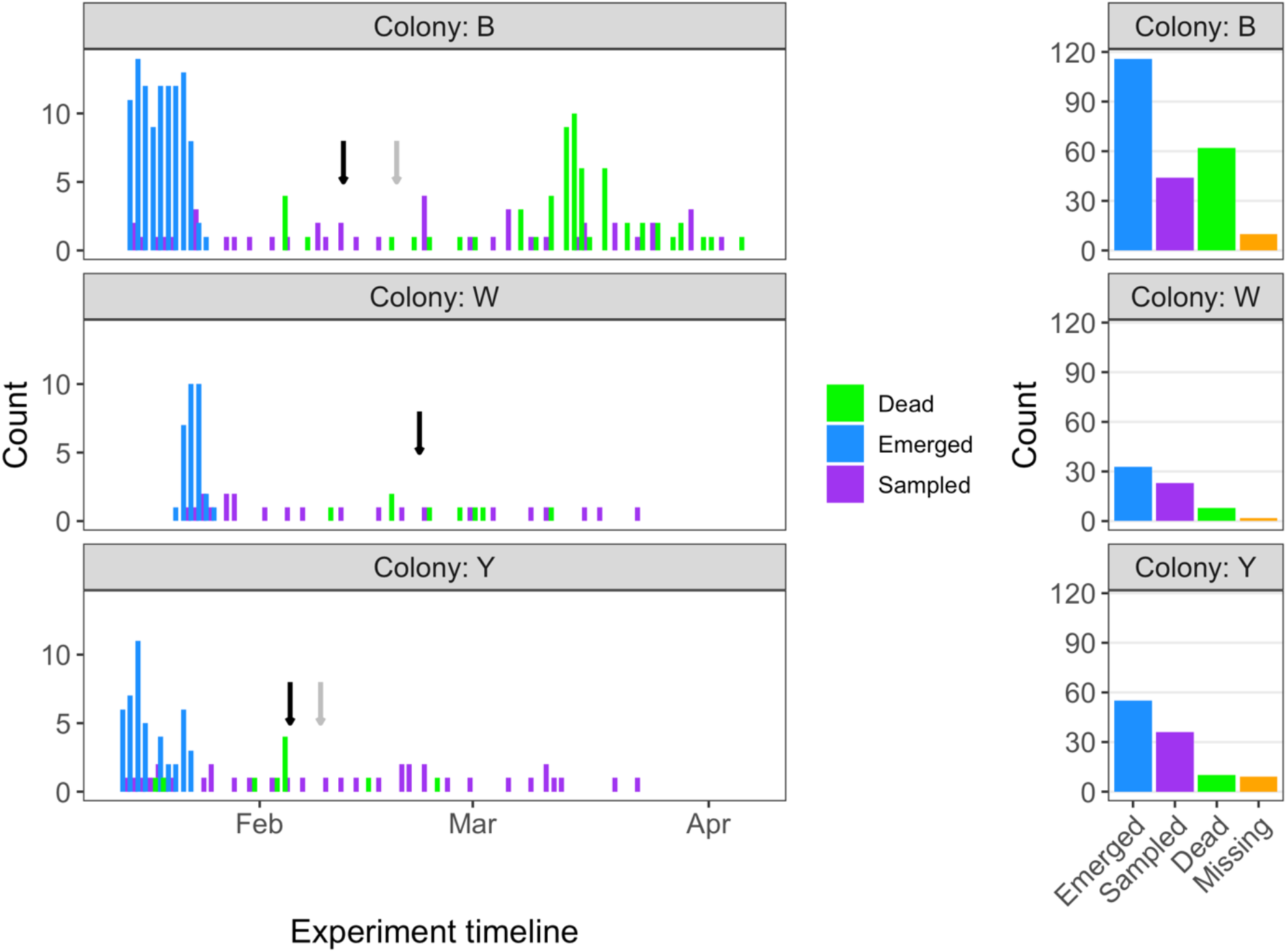
Timeline of worker emergence and tagging, gut sampling, and natural mortality. Sample sizes differed between colonies due to pre-existing variation in the number of brood. Total workers sampled were 44, 23, and 36 for Colonies B, W, and Y, respectively. Arrows indicate the first observation of reproductive bees in each colony (grey = queens, black = males); colony W did not produce queens. A small number of tagged bees were not recovered (right panel); these may have lost their tags, or died in a hidden part of the nest. Sampling continued until there were no live tagged bees left in the colony.

**Figure S2.**
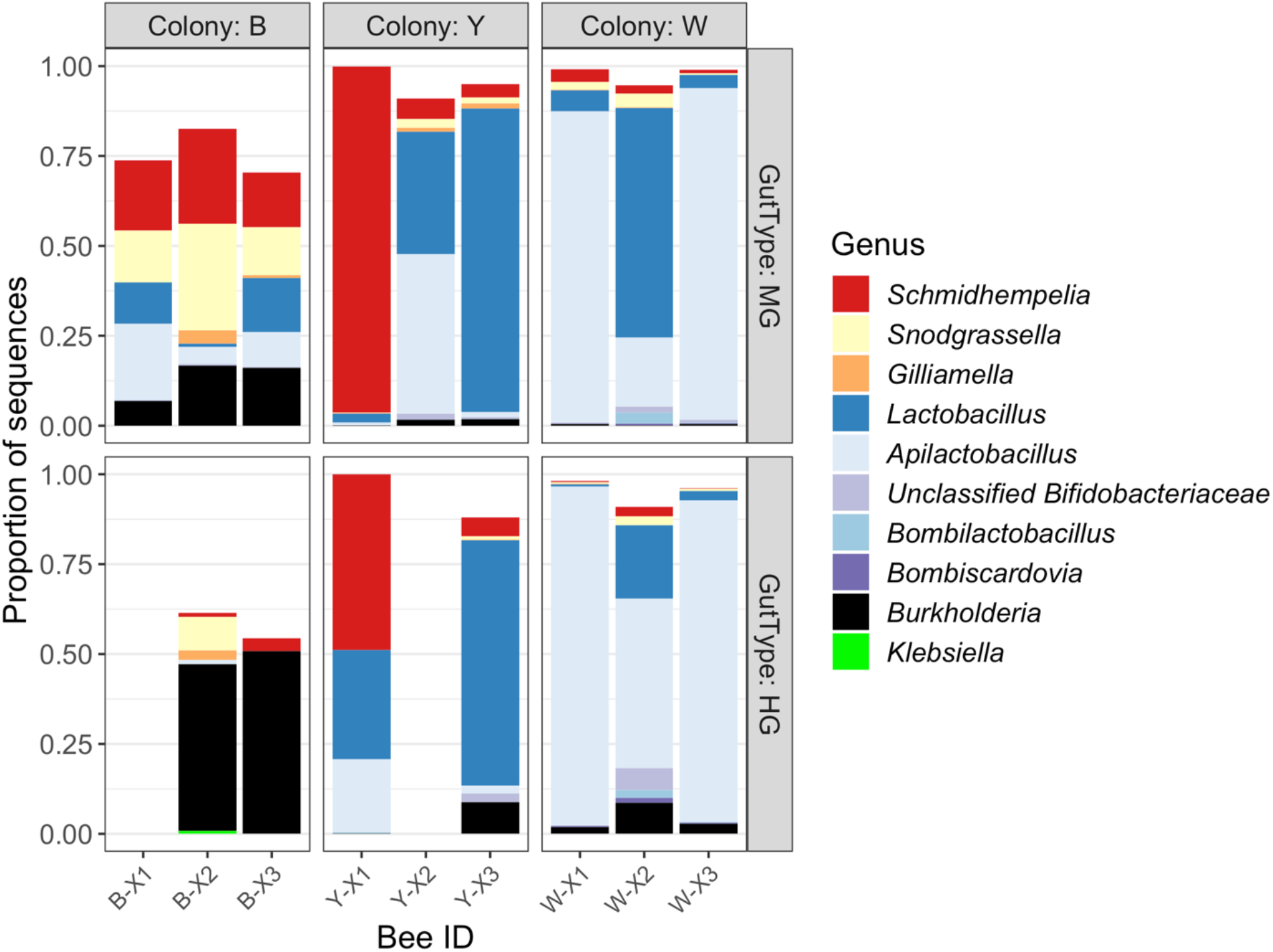
16S amplicon-based community composition in newly emerged bees, for both midgut (top row) and hindgut (bottom row) samples. Two samples were filtered out due to very low sequencing depth, probably as a result of low amounts of bacterial DNA (see Fig. 1A). *Burkholderia* is the most common taxon in our extraction blanks, and is likely to be a reagent contaminant in these gut samples.

**Figure S3.**
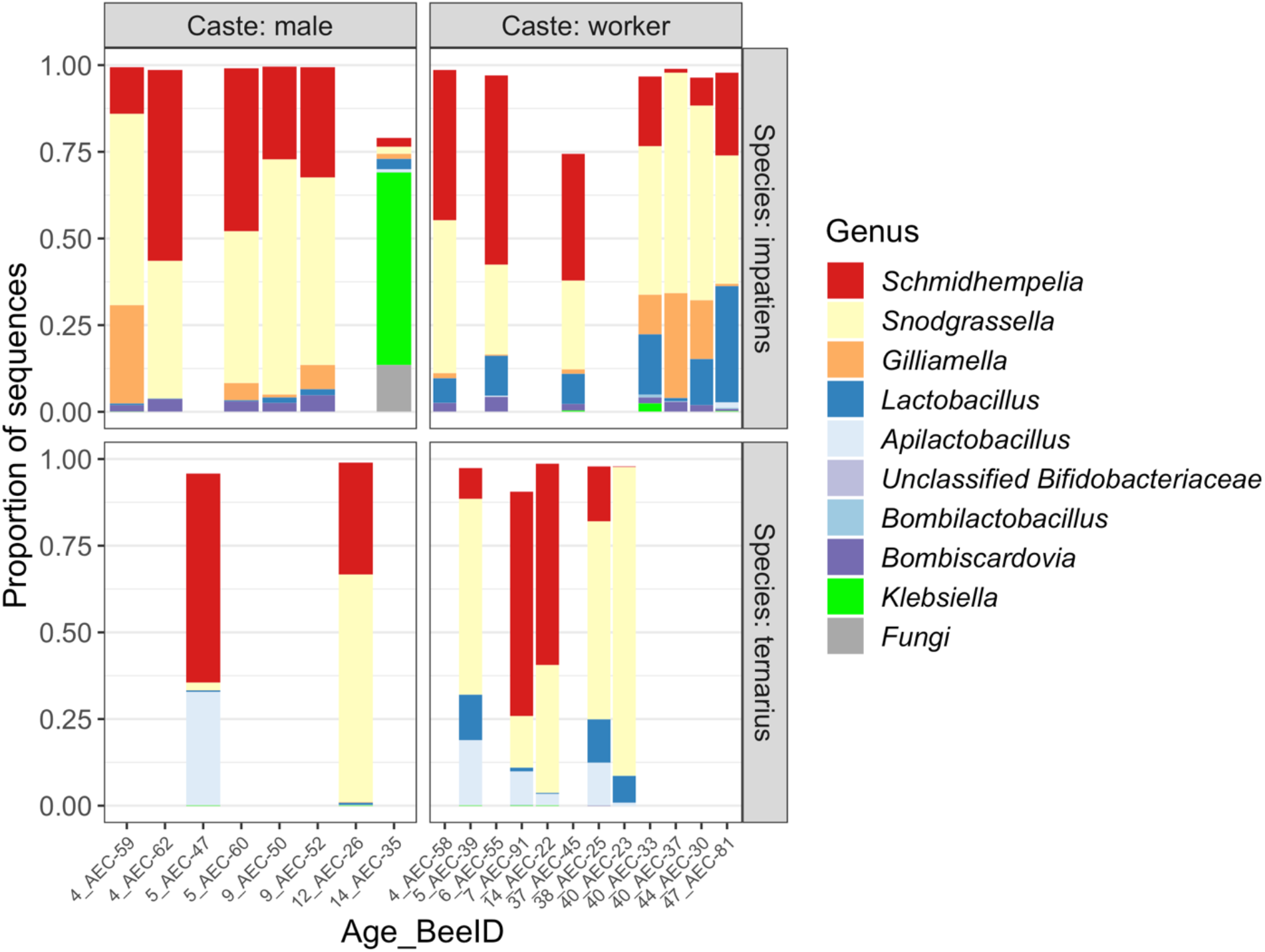
16S amplicon-based community composition of males and workers of two *Bombus* species reared indoors from wild queens. The age of each bee in days is noted by the number preceding the sample ID on the x axis. Fungi were identified based on an ASV matching the mitochondrial 16S rRNA gene of *Aspergillus* and *Mucor* in NCBI.

**Figure S4.**
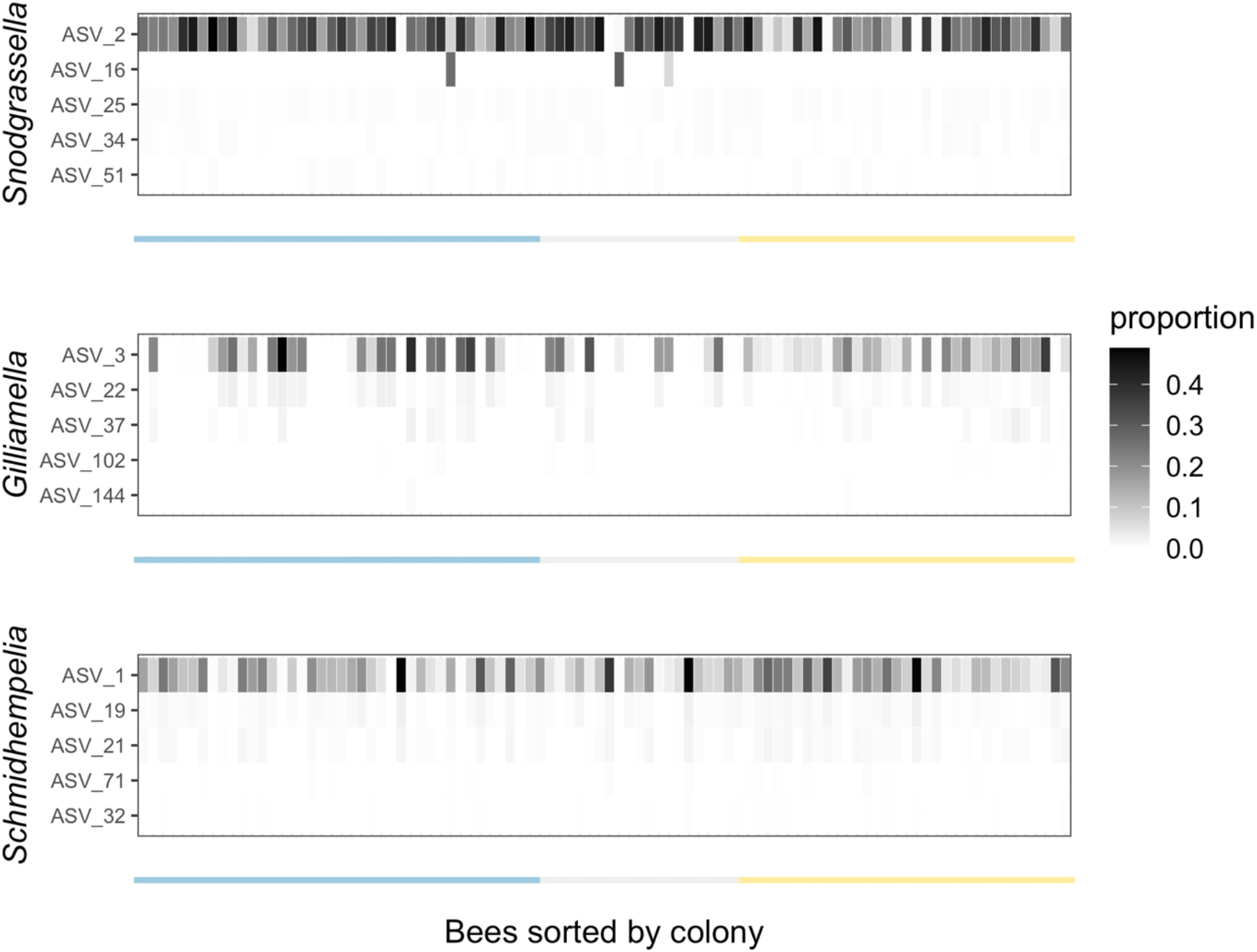
The existence of co-occurring subspecies (Fig. 3) and strain partioning among colonies (Fig. 4) is masked in 16S amplicon data. Here, distributions of the top five most abundant ASVs for the three most abundant core bacterial species are shown. Each column is an individual bee sorted by its colony affiliation.

**Figure S5.**
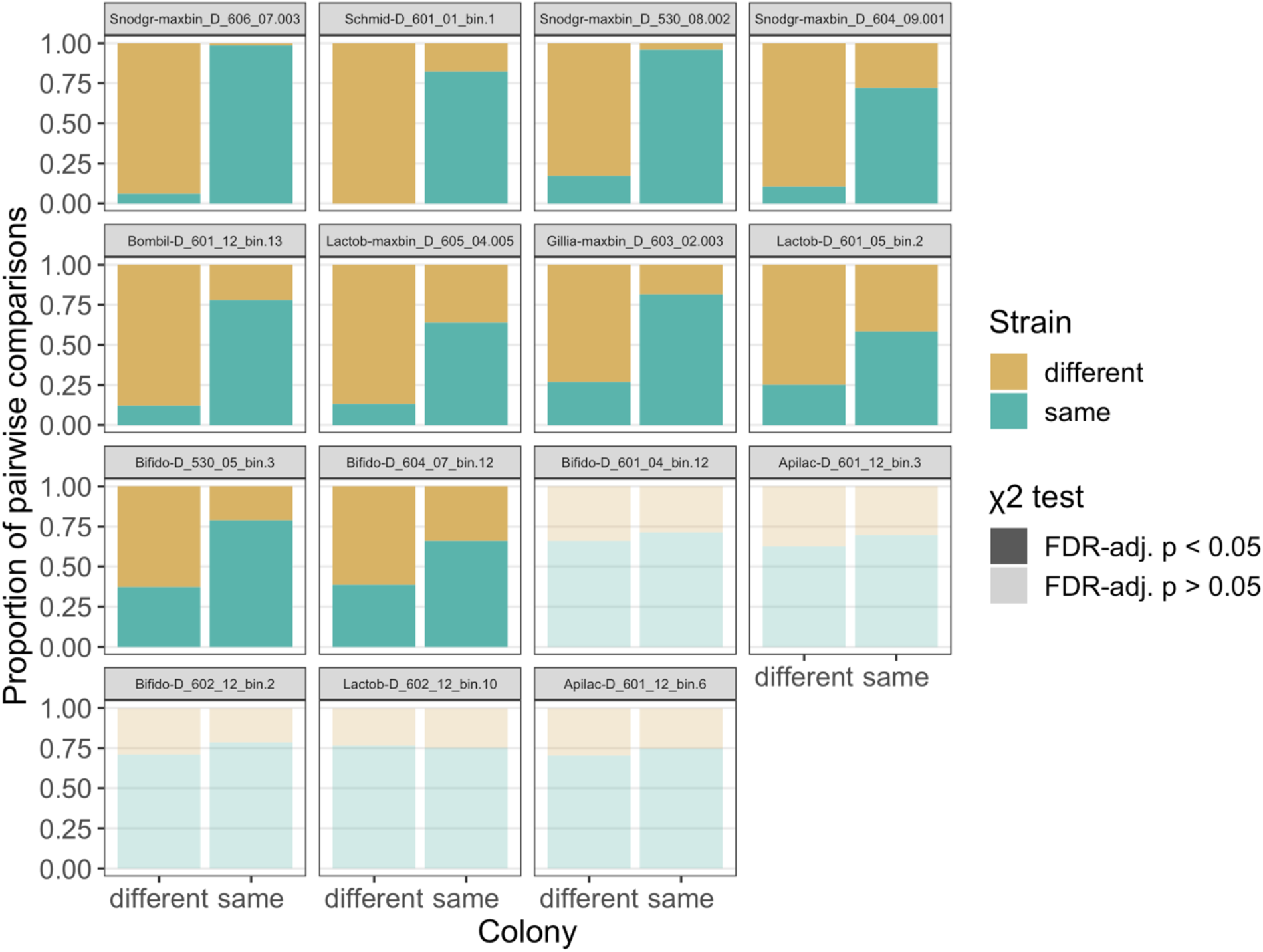
Strain-level colony partitioning broken down by MAG (Table S1). MAGs are sorted by the degree of colony differentiation; MAGs that did not exhibit a significant association with colony are translucent. *Snodgrassella, Schmidhempelia*, and *Gilliamella* are Gram-negative taxa, while the others are Gram-positive.

**Figure S6.**
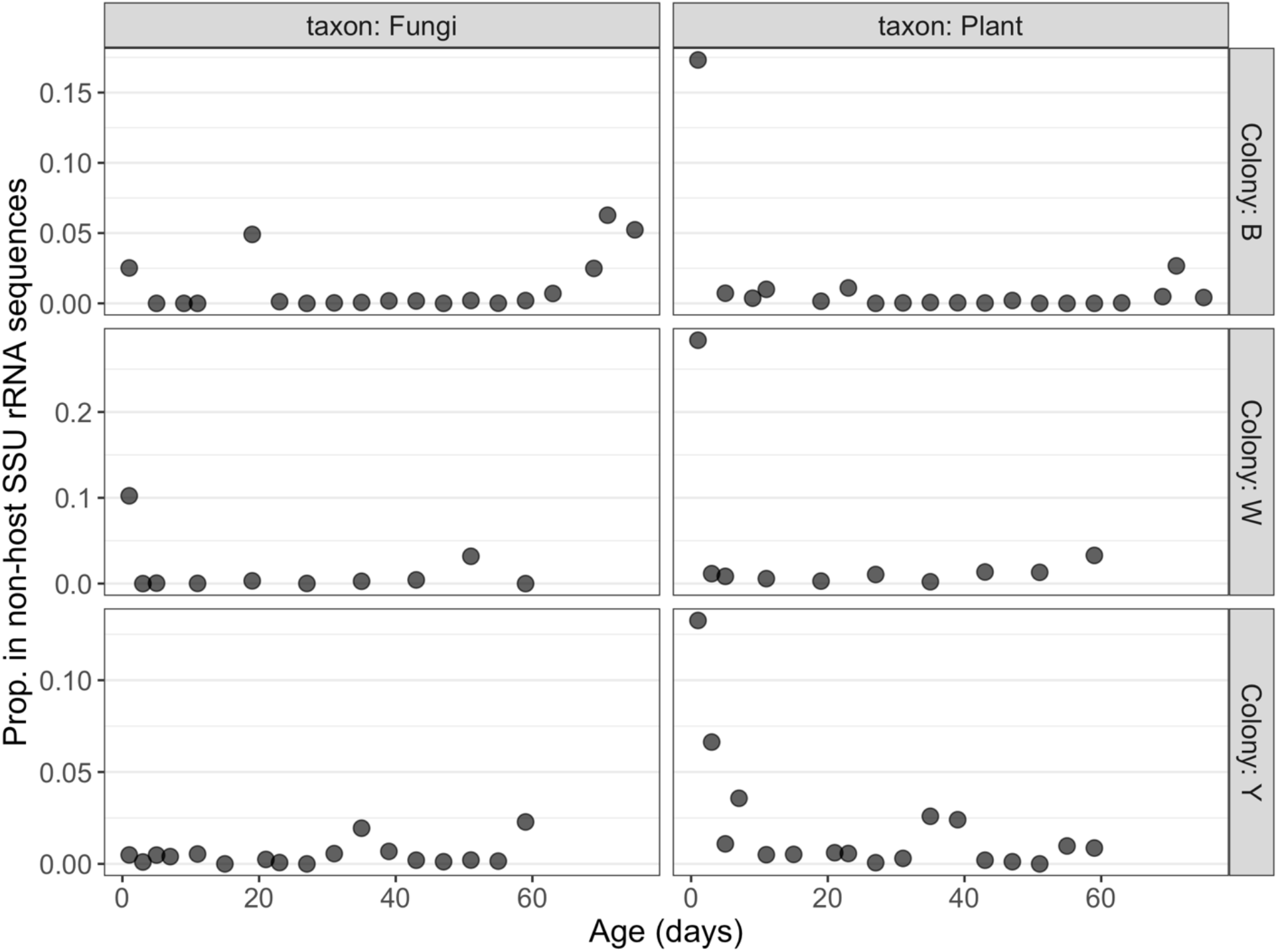
Composition of non-host, non-bacterial reads in 46 worker hindgut metagenomes using SSU rRNA genes identified by phyloFlash. The vast majority (median 99.12%) of non-host reads are bacterial.

**Figure S7.**
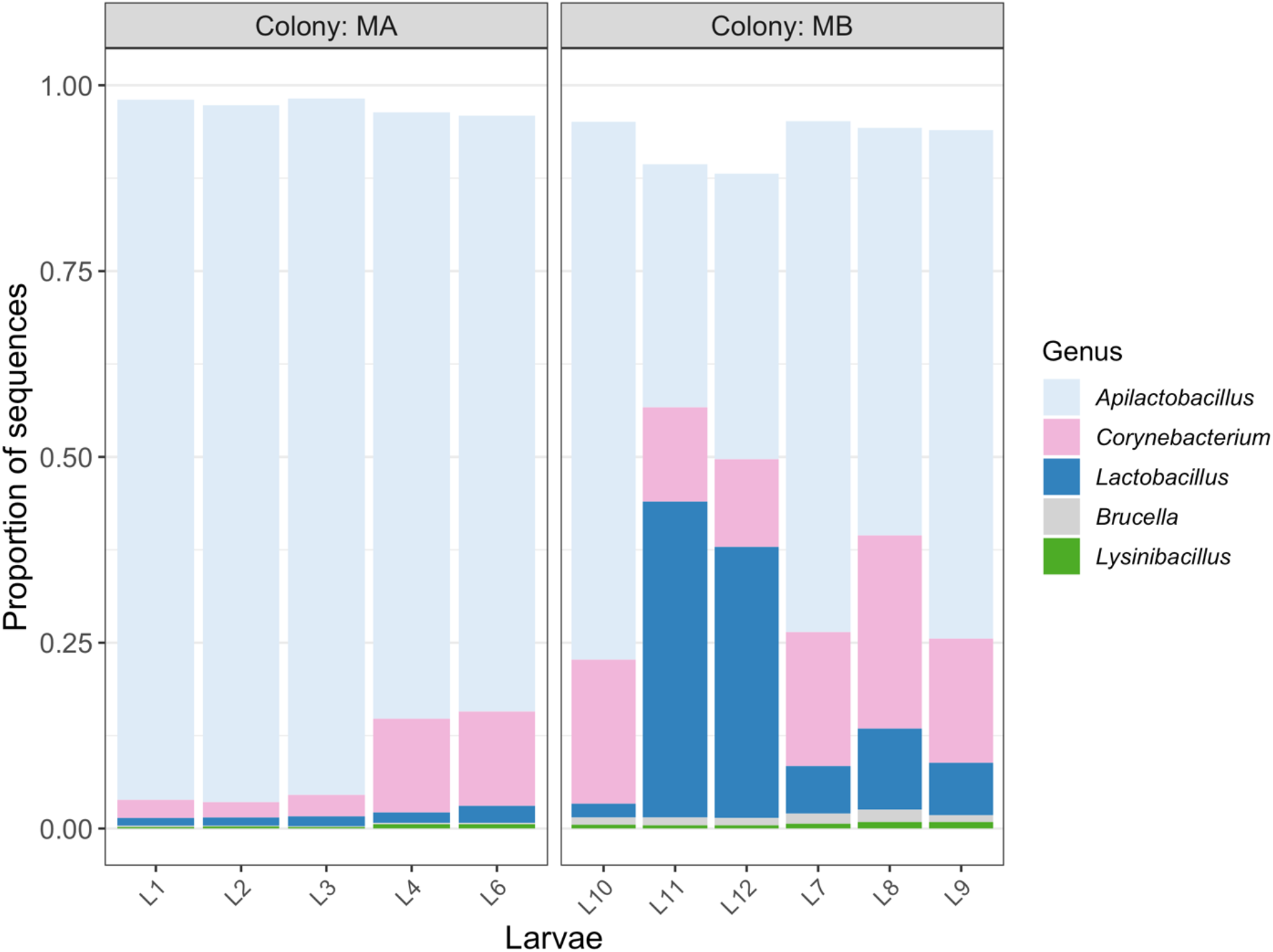
16S amplicon-based community composition in whole *Bombus impatiens* larvae from two replicate colonies. These colonies were obtained from the same commercial supplier as the colonies used for adult sampling. Aside from *Apilactobacillus* and *Lactobacillus*, taxa dominant in adult guts (Fig. 2A) were very rare or absent in these samples.

**Figure S8.**
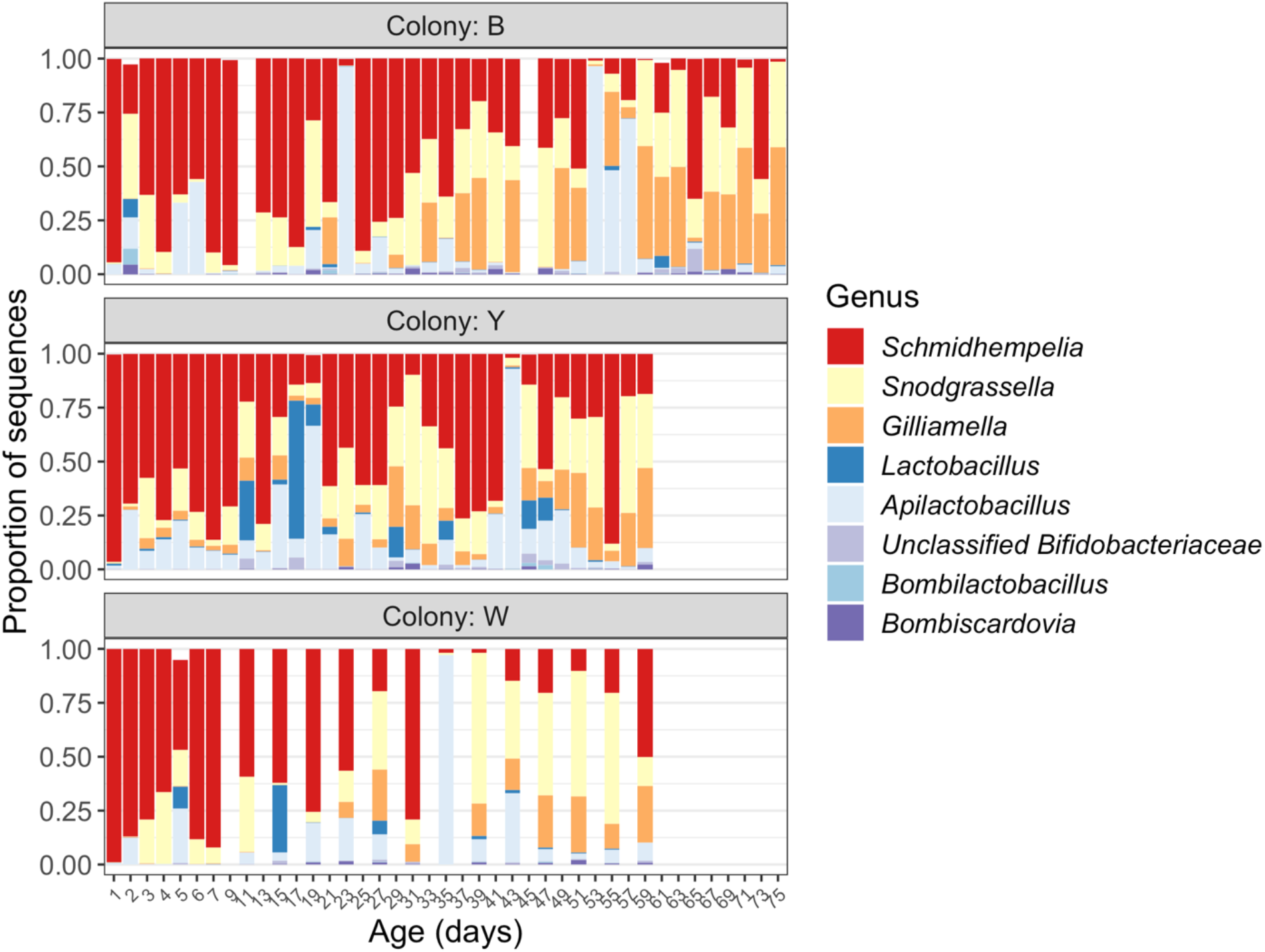
16S amplicon-based relative abundances of the top genus-level bacterial taxa in midguts of commercial *Bombus impatiens* workers. A total of 92 bees are shown, consisting of 39, 33, and 20 from colonies B, Y, and W, respectively. Two samples from colony B were filtered out due to low sequencing depth.

## References

1. Shade A, Caporaso JG, Handelsman J, Knight R, Fierer N. 2013. A meta-analysis of changes in bacterial and archaeal communities with time. ISME J 7:1493–506.

2. Nemergut DR, Schmidt SK, Fukami T, O’Neill SP, Bilinski TM, Stanish LF, Knelman JE, Darcy JL, Lynch RC, Wickey P, Ferrenberg S. 2013. Patterns and Processes of Microbial Community Assembly. Microbiology and Molecular Biology Reviews 77:342–356.

3. Fierer N, Nemergut D, Knight R, Craine JM. 2010. Changes through time: integrating microorganisms into the study of succession. Research in Microbiology 161:635–642.

4. Koenig JE, Spor A, Scalfone N, Fricker AD, Stombaugh J, Knight R, Angenent LT, Ley RE. 2011. Succession of microbial consortia in the developing infant gut microbiome. Proceedings of the National Academy of Sciences 108:4578–4585.

5. Hammer TJ, Moran NA. 2019. Links between metamorphosis and symbiosis in holometabolous insects. Phil Trans R Soc B 374:20190068.

6. Redford AJ, Fierer N. 2009. Bacterial succession on the leaf surface: a novel system for studying successional dynamics. Microbial Ecology 58:189–98.

7. Lovat LB. 1996. Age related changes in gut physiology and nutritional status. Gut 38:306– 309.

8. Rera M, Clark RI, Walker DW. 2012. Intestinal barrier dysfunction links metabolic and inflammatory markers of aging to death in Drosophila. Proceedings of the National Academy of Sciences 109:21528–21533.

9. Coon KL, Valzania L, McKinney DA, Vogel KJ, Brown MR, Strand MR. 2017. Bacteria-mediated hypoxia functions as a signal for mosquito development. Proceedings of the National Academy of Sciences 114:E5362–E5369.

10. Heintz C, Mair W. 2014. You Are What You Host: Microbiome Modulation of the Aging Process. Cell 156:4.

11. Koropatnick TA, Engle JT, Apicella MA, Stabb EV, Goldman WE, McFall-Ngai MJ. 2004. Microbial Factor-Mediated Development in a Host-Bacterial Mutualism. Science 306:1186–1188.

12. Gilbert JA, Blaser MJ, Caporaso JG, Jansson JK, Lynch SV, Knight R. 2018. Current understanding of the human microbiome. Nat Med 24:392–400.

13. Vega NM, Gore J. 2017. Stochastic assembly produces heterogeneous communities in the Caenorhabditis elegans intestine. PLoS Biol 15:e2000633.

14. Obadia B, Güvener ZT, Zhang V, Ceja-Navarro JA, Brodie EL, Ja WW, Ludington WB. 2017. Probabilistic Invasion Underlies Natural Gut Microbiome Stability. Current Biology 27:1999-2006.e8.

15. Ellegaard KM, Engel P. 2019. Genomic diversity landscape of the honey bee gut microbiota. Nature Communications 10:446.

16. Costello EK, Lauber CL, Hamady M, Fierer N, Gordon JI, Knight R. 2009. Bacterial community variation in human body habitats across space and time. Science 326:1694– 1697.

17. Stephens WZ, Burns AR, Stagaman K, Wong S, Rawls JF, Guillemin K, Bohannan BJM. 2016. The composition of the zebrafish intestinal microbial community varies across development. ISME J 10:644–654.

18. Laughton AM, Fan MH, Gerardo NM. 2014. The Combined Effects of Bacterial Symbionts and Aging on Life History Traits in the Pea Aphid, Acyrthosiphon pisum. Appl Environ Microbiol 80:470–477.

19. Jones EW, Carlson JM, Sivak DA, Ludington WB. 2022. Stochastic microbiome assembly depends on context. Proc Natl Acad Sci USA 119:e2115877119.

20. Bäckhed F, Roswall J, Peng Y, Feng Q, Jia H, Kovatcheva-Datchary P, Li Y, Xia Y, Xie H, Zhong H, Khan MT, Zhang J, Li J, Xiao L, Al-Aama J, Zhang D, Lee YS, Kotowska D, Colding C, Tremaroli V, Yin Y, Bergman S, Xu X, Madsen L, Kristiansen K, Dahlgren J, Wang J. 2015. Dynamics and Stabilization of the Human Gut Microbiome during the First Year of Life. Cell Host & Microbe 17:690–703.

21. Stewart CJ, Ajami NJ, O’Brien JL, Hutchinson DS, Smith DP, Wong MC, Ross MC, Lloyd RE, Doddapaneni H, Metcalf GA, Muzny D, Gibbs RA, Vatanen T, Huttenhower C, Xavier RJ, Rewers M, Hagopian W, Toppari J, Ziegler A-G, She J-X, Akolkar B, Lernmark A, Hyoty H, Vehik K, Krischer JP, Petrosino JF. 2018. Temporal development of the gut microbiome in early childhood from the TEDDY study. Nature 562:583–588.

22. Powell JE, Martinson VG, Urban-Mead K, Moran NA. 2014. Routes of acquisition of the gut microbiota of the honey bee Apis mellifera. Appl Environ Microbiol 80:7378–7387.

23. Claesson MJ, Jeffery IB, Conde S, Power SE, O’Connor EM, Cusack S, Harris HMB, Coakley M, Lakshminarayanan B, O’Sullivan O, Fitzgerald GF, Deane J, O’Connor M, Harnedy N, O’Connor K, O’Mahony D, van Sinderen D, Wallace M, Brennan L, Stanton C, Marchesi JR, Fitzgerald AP, Shanahan F, Hill C, Ross RP, O’Toole PW. 2012. Gut microbiota composition correlates with diet and health in the elderly. Nature 488:178–184.

24. Claesson MJ, Cusack S, O’Sullivan O, Greene-Diniz R, de Weerd H, Flannery E, Marchesi JR, Falush D, Dinan T, Fitzgerald G, Stanton C, van Sinderen D, O’Connor M, Harnedy N, O’Connor K, Henry C, O’Mahony D, Fitzgerald AP, Shanahan F, Twomey C, Hill C, Ross RP, O’Toole PW. 2011. Composition, variability, and temporal stability of the intestinal microbiota of the elderly. Proceedings of the National Academy of Sciences of the United States of America 108:4586–91.

25. Wilmanski T, Diener C, Rappaport N, Patwardhan S, Wiedrick J, Lapidus J, Earls JC, Zimmer A, Glusman G, Robinson M, Yurkovich JT, Kado DM, Cauley JA, Zmuda J, Lane NE, Magis AT, Lovejoy JC, Hood L, Gibbons SM, Orwoll ES, Price ND. 2021. Gut microbiome pattern reflects healthy ageing and predicts survival in humans. Nat Metab 3:274–286.

26. Biagi E, Nylund L, Candela M, Ostan R, Bucci L, Pini E, Nikkïla J, Monti D, Satokari R, Franceschi C, Brigidi P, De Vos W. 2010. Through Ageing, and Beyond: Gut Microbiota and Inflammatory Status in Seniors and Centenarians. PLoS ONE 5:e10667.

27. Smith P, Willemsen D, Popkes M, Metge F, Gandiwa E, Reichard M, Valenzano DR. 2017. Regulation of life span by the gut microbiota in the short-lived African turquoise killifish. eLife 6:e27014.

28. Thevaranjan N, Puchta A, Schulz C, Naidoo A, Szamosi JC, Verschoor CP, Loukov D, Schenck LP, Jury J, Foley KP, Schertzer JD, Larché MJ, Davidson DJ, Verdú EF, Surette MG, Bowdish DME. 2017. Age-Associated Microbial Dysbiosis Promotes Intestinal Permeability, Systemic Inflammation, and Macrophage Dysfunction. Cell Host & Microbe 21:455–466.

29. Erkosar B, Leulier F. 2014. Transient adult microbiota, gut homeostasis and longevity: Novel insights from the Drosophila model. FEBS Letters 588:4250–4257.

30. Clark RI, Salazar A, Yamada R, Fitz-Gibbon S, Morselli M, Alcaraz J, Rana A, Rera M, Pellegrini M, Ja WW, Walker DW. 2015. Distinct Shifts in Microbiota Composition during Drosophila Aging Impair Intestinal Function and Drive Mortality. Cell Reports 12:1656– 1667.

31. Fransen F, van Beek AA, Borghuis T, Aidy SE, Hugenholtz F, van der Gaast – de Jongh C, Savelkoul HFJ, De Jonge MI, Boekschoten MV, Smidt H, Faas MM, de Vos P. 2017. Aged Gut Microbiota Contributes to Systemical Inflammaging after Transfer to Germ-Free Mice. Front Immunol 8:1385.

32. Reese AT, Phillips SR, Owens LA, Venable EM, Langergraber KE, Machanda ZP, Mitani JC, Muller MN, Watts DP, Wrangham RW, Goldberg TL, Emery Thompson M, Carmody RN. 2021. Age Patterning in Wild Chimpanzee Gut Microbiota Diversity Reveals Differences from Humans in Early Life. Current Biology 31:613–620.

33. Risely A, Wilhelm K, Clutton-Brock T, Manser MB, Sommer S. 2021. Diurnal oscillations in gut bacterial load and composition eclipse seasonal and lifetime dynamics in wild meerkats. Nat Commun 12:6017.

34. Kwong WK, Moran NA. 2016. Gut microbial communities of social bees. Nat Rev Microbiol 14:374–384.

35. Hammer TJ, L. E, Martin AN, Moran NA. 2021. The gut microbiota of bumblebees. Insect Soc 68:287–301.

36. Menezes C, Vollet-Neto A, Marsaioli AJ, Zampieri D, Fontoura IC, Luchessi AD, Imperatriz-Fonseca VL. 2015. A Brazilian Social Bee Must Cultivate Fungus to Survive. Current Biology 25:2851–2855.

37. Goulson D, Nicholls E, Botias C, Rotheray EL. 2015. Bee declines driven by combined stress from parasites, pesticides, and lack of flowers. Science 347:1255957.

38. Faust K, Lahti L, Gonze D, de Vos WM, Raes J. 2015. Metagenomics meets time series analysis: unraveling microbial community dynamics. Current Opinion in Microbiology 25:56–66.

39. Zaneveld JR, McMinds R, Vega Thurber R. 2017. Stress and stability: applying the Anna Karenina principle to animal microbiomes. Nature Microbiology 2:17121.

40. Cerqueira AES, Hammer TJ, Moran NA, Santana WC, Kasuya MCM, da Silva CC. 2021. Extinction of anciently associated gut bacterial symbionts in a clade of stingless bees. ISME J 15:2813–2816.

41. Kwong WK, Medina LA, Koch H, Sing K-W, Soh EJY, Ascher JS, Jaffé R, Moran NA. 2017. Dynamic microbiome evolution in social bees. Sci Adv 3:e1600513.

42. Grime JP. 1974. Vegetation classification by reference to strategies. Nature 250:26–31.

43. Ho A, Lonardo DPD, Bodelier PLE. 2017. Revisiting life strategy concepts in environmental microbial ecology. Microbiology Ecology 93:fix006.

44. Martinson VG, Moy J, Moran NA. 2012. Establishment of characteristic gut bacteria during development of the honeybee worker. Appl Environ Microbiol 78:2830–2840.

45. Kešnerová L, Emery O, Troilo M, Liberti J, Erkosar B, Engel P. 2020. Gut microbiota structure differs between honeybees in winter and summer. ISME J 14:801–814.

46. Maes PW, Floyd AS, Mott BM, Anderson KE. 2021. Overwintering Honey Bee Colonies: Effect of Worker Age and Climate on the Hindgut Microbiota. Insects 12:224.

47. Kapheim KM, Rao VD, Yeoman CJ, Wilson BA, White BA, Goldenfeld N, Robinson GE. 2015. Caste-Specific Differences in Hindgut Microbial Communities of Honey Bees (Apis mellifera). PLOS ONE 10:e0123911.

48. Seeley T. 1982. Adaptive significance of the age polyethism schedule in honeybee colonies. Behavioral Ecology and Sociobiology 11:287–293.

49. Winston M. 1991. The biology of the honey bee. Harvard University Press, Cambridge, Massachusetts.

50. Steinmann N, Corona M, Neumann P, Dainat B. 2015. Overwintering Is Associated with Reduced Expression of Immune Genes and Higher Susceptibility to Virus Infection in Honey Bees. PLoS ONE 10:e0129956.

51. Amdam GV, Omholt SW. 2002. The Regulatory Anatomy of Honeybee Lifespan. Journal of Theoretical Biology 216:209–228.

52. Cameron SA. 1989. Temporal patterns of division of labor among workers in the primitively eusocial bumble bee, Bombus griseocollis (Hymenoptera: Apidae). Ethology 80:137–151.

53. Martinson VG, Magoc T, Koch H, Salzberg SL, Moran NA. 2014. Genomic features of a bumble bee symbiont reflect its host environment. Appl Environ Microbiol 80:3793–3803.

54. Li J, Powell JE, Guo J, Evans JD, Wu J, Williams P, Lin Q, Moran NA, Zhang Z. 2015. Two gut community enterotypes recur in diverse bumblebee species. Curr Biol 25:R652– R653.

55. Koch H, Cisarovsky G, Schmid-Hempel P. 2012. Ecological effects on gut bacterial communities in wild bumblebee colonies. J Anim Ecol 81:1202–1210.

56. Su Q, Wang Q, Mu X, Chen H, Meng Y, Zhang X, Zheng L, Hu X, Zhai Y, Zheng H. 2021. Strain-level analysis reveals the vertical microbial transmission during the life cycle of bumblebee. Microbiome 9:216.

57. Wang L, Wu J, Li K, Sadd BM, Guo Y, Zhuang D, Zhang Z, Chen Y, Evans JD, Guo J, Zhang Z, Li J. 2019. Dynamic changes of gut microbial communities of bumble bee queens through important life stages. mSystems 4:e00631–19.

58. Meeus I, Mommaerts V, Billiet A, Mosallanejad H, Van De Wiele T, Wäckers F, Smagghe G. 2013. Assessment of mutualism between Bombus terrestris and its microbiota by use of microcolonies. Apidologie 44:708–719.

59. Evans JD, Lopez DL. 2004. Bacterial Probiotics Induce an Immune Response in the Honey Bee (Hymenoptera: Apidae). Journal of Economic Entomology 97:752–756.

60. Kwong WK, Mancenido AL, Moran NA. 2017. Immune system stimulation by the native gut microbiota of honey bees. R Soc Open Sci 4:170003.

61. Peters A, Delhey K, Nakagawa S, Aulsebrook A, Verhulst S. 2019. Immunosenescence in wild animals: meta-analysis and outlook. Ecol Lett 22:1709–1722.

62. Heinze J, Giehr J. 2021. The plasticity of lifespan in social insects. Phil Trans R Soc B 376:20190734.

63. Keller L, Genoud M. 1997. Extraordinary lifespans in ants: a test of evolutionary theories of ageing. Nature 389:958–960.

64. Goulson D. 2003. Bumblebees: Behaviour and Ecology. Oxford University Press.

65. Muller CB, Schmid-Hempel P. 1992. Variation in Life-History Pattern in Relation to Worker Mortality in the Bumble-Bee, Bombus lucorum. Functional Ecology 6:48.

66. Goldblatt JW, Fell RD. 1987. Adult longevity of workers of the bumble bees Bombus fervidus (F.) and Bombus pennsylvanicus (De Geer) (Hymenoptera: Apidae). Can J Zool 65:2349–2353.

67. Moret Y, Schmid-Hempel P. 2009. Immune responses of bumblebee workers as a function of individual and colony age: senescence versus plastic adjustment of the immune function. Oikos 118:371–378.

68. Salter SJ, Cox MJ, Turek EM, Calus ST, Cookson WO, Moffatt MF, Turner P, Parkhill J, Loman NJ, Walker AW. 2014. Reagent and laboratory contamination can critically impact sequence-based microbiome analyses. BMC Biology 12:87.

69. Meeus I, Parmentier L, Billiet A, Maebe K, Van Nieuwerburgh F, Deforce D, Wäckers F, Vandamme P, Smagghe G. 2015. 16S rRNA amplicon sequencing demonstrates that indoor-reared bumblebees (Bombus terrestris) harbor a core subset of bacteria normally associated with the wild host. PLoS One 10:1–15.

70. Parmentier L, Meeus I, Mosallanejad H, de Graaf DC, Smagghe G. 2016. Plasticity in the gut microbial community and uptake of Enterobacteriaceae (Gammaproteobacteria) in Bombus terrestris bumblebees’ nests when reared indoors and moved to an outdoor environment. Apidologie 47:237–250.

71. Krams R, Gudra D, Popovs S, Willow J, Krama T, Munkevics M, Megnis K, Jõers P, Fridmanis D, Contreras Garduño J, Krams IA. 2022. Dominance of Fructose-Associated Fructobacillus in the Gut Microbiome of Bumblebees (Bombus terrestris) Inhabiting Natural Forest Meadows. Insects 13:98.

72. Bosmans L, Pozo MI, Verreth C, Crauwels S, Wilberts L, Sobhy IS, Wäckers F, Jacquemyn H, Lievens B. 2018. Habitat-specific variation in gut microbial communities and pathogen prevalence in bumblebee queens (Bombus terrestris). PLoS One 13:e0204612.

73. Olm MR, Crits-Christoph A, Bouma-Gregson K, Firek BA, Morowitz MJ, Banfield JF. 2021. inStrain profiles population microdiversity from metagenomic data and sensitively detects shared microbial strains. Nat Biotechnol 39:727–736.

74. Lemaitre B, Hoffmann J. 2007. The Host Defense of Drosophila melanogaster. Annu Rev Immunol 25:697–743.

75. Engel P, Moran NA. 2013. The gut microbiota of insects – diversity in structure and function. FEMS Microbiol Rev 37:699–735.

76. Ha E-M, Oh C-T, Ryu J-H, Bae Y-S, Kang S-W, Jang I, Brey PT, Lee W-J. 2005. An Antioxidant System Required for Host Protection against Gut Infection in Drosophila. Developmental Cell 8:125–132.

77. Sauers LA, Sadd BM. 2019. An interaction between host and microbe genotypes determines colonization success of a key bumble bee gut microbiota member. Evolution 73:2333–2342.

78. Hammer TJ, L. E, Moran NA. 2021. Thermal niches of specialized gut symbionts: the case of social bees. Proc R Soc B 288:20201480.

79. Koch H, Schmid-Hempel P. 2011. Socially transmitted gut microbiota protect bumble bees against an intestinal parasite. PNAS 108:19288–19292.

80. Raymann K, Shaffer Z, Moran NA. 2017. Antibiotic exposure perturbs the gut microbiota and elevates mortality in honeybees. PLoS Biol 15:e2001861.

81. Sanidad KZ, Zeng MY. 2020. Neonatal gut microbiome and immunity. Current Opinion in Microbiology 56:30–37.

82. Rao C, Coyte KZ, Bainter W, Geha RS, Martin CR, Rakoff-Nahoum S. 2021. Multi-kingdom ecological drivers of microbiota assembly in preterm infants. Nature 591:633– 638.

83. Bokulich NA, Chung J, Battaglia T, Henderson N, Jay M, Li H, D. Lieber A, Wu F, Perez-Perez GI, Chen Y, Schweizer W, Zheng X, Contreras M, Dominguez-Bello MG, Blaser MJ. 2016. Antibiotics, birth mode, and diet shape microbiome maturation during early life. Sci Transl Med 8:343ra82.

84. Moran NA, Ochman H, Hammer TJ. 2019. Evolutionary and ecological consequences of gut microbial communities. Annu Rev Ecol Evol Syst 50:451–475.

85. Tung J, Barreiro LB, Burns MB, Grenier JC, Lynch J, Grieneisen LE, Altmann J, Alberts SC, Blekhman R, Archie EA. 2015. Social networks predict gut microbiome composition in wild baboons. eLife 2015:1–18.

86. Moeller AH, Suzuki TA, Phifer-Rixey M, Nachman MW. 2018. Transmission modes of the mammalian gut microbiota. Science 362:453–457.

87. Seeley T, Heinrich B. 1981. Regulation of temperature in nests of social insects, p. 159– 234. In Insect Thermoregulation. Wiley, New York.

88. Conwill A, Kuan AC, Damerla R, Tripp AD, Alm EJ, Lieberman TD. 2022. Anatomy promotes neutral coexistence of strains in the human skin microbiome. Cell Host & Microbe 30:171–182.

89. Livingston G, Matias M, Calcagno V, Barbera C, Combe M, Leibold MA, Mouquet N. 2012. Competition–colonization dynamics in experimental bacterial metacommunities. Nat Commun 3:1234.

90. Yawata Y, Cordero OX, Menolascina F, Hehemann J-H, Polz MF, Stocker R. 2014. Competition–dispersal tradeoff ecologically differentiates recently speciated marine bacterioplankton populations. Proc Natl Acad Sci USA 111:5622–5627.

91. Smith GR, Steidinger BS, Bruns TD, Peay KG. 2018. Competition-colonization tradeoffs structure fungal diversity. ISME Journal 12:1758–1767.

92. Steele MI, Kwong WK, Whiteley M, Moran NA. 2017. Diversification of type VI secretion system toxins reveals ancient antagonism among bee gut microbes. mBio 8:e01630–17.

93. Rothman J, Russell K, Leger L, McFrederick Q, Graystock P. 2020. The direct and indirect effects of environmental toxicants on the health of bumble bees and their microbiomes. Proc R Soc B 287:20200980.

94. Rothman JA, Leger L, Graystock P, Russell K, McFrederick QS. 2019. The bumble bee microbiome increases survival of bees exposed to selenate toxicity. Environ Microbiol 21:3417–3429.

95. Palmer-Young EC, Ngor L, Burciaga Nevarez R, Rothman JA, Raffel TR, McFrederick QS. 2019. Temperature dependence of parasitic infection and gut bacterial communities in bumble bees. Environ Microbiol 21:4706–4723.

96. Diez-Méndez D, Bodawatta KH, Freiberga I, Klečková I, Jønsson KA, Poulsen M, Sam K. 2022. Gut microbiome disturbances of altricial Blue and Great tit nestlings are countered by continuous microbial inoculations from parental microbiomes. bioRxiv https://doi.org/10.1101/2022.02.20.481211.

97. Yarlagadda K, Razik I, Malhi RS, Carter GG. 2021. Social convergence of gut microbiomes in vampire bats. Biol Lett 17:20210389.

98. Ellington CP, Machin KE, Casey TM. 1990. Oxygen consumption of bumblebees in forward flight. Nature 347:472–473.

99. Schmid-Hempel P, Wolf T. 1988. Foraging Effort and Life Span of Workers in a Social Insect. The Journal of Animal Ecology 57:509.

100. Doums C, Schmid-Hempel P. 2000. Immunocompetence in workers of a social insect, Bombus terrestris L., in relation to foraging activity and parasitic infection. Canadian Journal of Zoology 78:1060–1066.

101. Kirkwood T, Rose M. 1991. Evolution of senescence: late survival sacrificed for reproduction. Phil Trans R Soc Lond B 332:15–24.

102. Williams G. 1957. Pleiotropy, natural selection, and the evolution of senescence. Evolution 11:398–411.

103. Hamilton WD. 1966. The moulding of senescence by natural selection. Journal of Theoretical Biology 12:12–45.

104. Jones OR, Scheuerlein A, Salguero-Gómez R, Camarda CG, Schaible R, Casper BB, Dahlgren JP, Ehrlén J, García MB, Menges ES, Quintana-Ascencio PF, Caswell H, Baudisch A, Vaupel JW. 2014. Diversity of ageing across the tree of life. Nature 505:169– 173.

105. Hagbery J, Nieh JC. 2012. Individual lifetime pollen and nectar foraging preferences in bumble bees. Naturwissenschaften 99:821–832.

106. Kelemen EP, Cao N, Cao T, Davidowitz G, Dornhaus A. 2019. Metabolic rate predicts the lifespan of workers in the bumble bee Bombus impatiens. Apidologie 50:195–203.

107. Blacher P, Huggins TJ, Bourke AFG. 2017. Evolution of ageing, costs of reproduction and the fecundity–longevity trade-off in eusocial insects. Proc R Soc B 284:20170380.

108. Smeets P, Duchateau MJ. 2003. Longevity of Bombus terrestris workers (Hymenoptera: Apidae) in relation to pollen availability, in the absence of foraging. Apidologie 34:333– 337.

109. Brian AD. 1952. Division of Labour and Foraging in Bombus agrorum Fabricius. The Journal of Animal Ecology 21:223–240.

110. Cartar RV. 1992. Morphological Senescence and Longevity: An Experiment Relating Wing Wear and Life Span in Foraging Wild Bumble Bees. The Journal of Animal Ecology 61:225–231.

111. Rodd FH, Plowright RC, Owen RE. 1980. Mortality rates of adult bumble bee workers (Hymenoptera: Apidae). Can J Zool 58:1718–1721.

112. Powell JE, Carver Z, Leonard SP, Moran NA. 2021. Field-Realistic Tylosin Exposure Impacts Honey Bee Microbiota and Pathogen Susceptibility, Which Is Ameliorated by Native Gut Probiotics. Microbiol Spectr 9:e00103–21.

113. Martin M. 2011. Cutadapt removes adapter sequences from high-throughput sequencing reads. EMBnet.journal 17:10–12.

114. Callahan BJ, McMurdie PJ, Rosen MJ, Han AW, Johnson AJA, Holmes SP. 2016. DADA2: High-resolution sample inference from Illumina amplicon data. Nat Methods 13:581–583.

115. Quast C, Pruesse E, Yilmaz P, Gerken J, Schweer T, Yarza P, Peplies J, Glöckner FO. 2013. The SILVA ribosomal RNA gene database project: Improved data processing and web-based tools. Nucleic Acids Research 41:590–596.

116. Hammer TJ, Dickerson JC, McMillan WO, Fierer N. 2020. Heliconius butterflies host characteristic and phylogenetically structured adult-stage microbiomes. Appl Environ Microbiol 86:e02007–20.

117. Sadd BM, et al. 2015. The genomes of two key bumblebee species with primitive eusocial organization. Genome Biol 16:76.

118. Dobin A, Davis CA, Schlesinger F, Drenkow J, Zaleski C, Jha S, Batut P, Chaisson M, Gingeras TR. 2013. STAR: ultrafast universal RNA-seq aligner. Bioinformatics 29:15–21.

119. Law C, Alhamdoosh M, Su S, Dong X, Tian L, Smyth G, Ritchie M. 2018. RNA-seq analysis is easy as 1-2-3 with limma, Glimma and edgeR. F1000Research 5:1408.

120. Ritchie ME, Phipson B, Wu D, Hu Y, Law CW, Shi W, Smyth GK. 2015. limma powers differential expression analyses for RNA-sequencing and microarray studies. Nucleic Acids Research 43:e47.

121. Robinson MD, McCarthy DJ, Smyth GK. 2010. edgeR: a Bioconductor package for differential expression analysis of digital gene expression data. Bioinformatics 26:139–140.

122. Langmead B, Salzberg SL. 2012. Fast gapped-read alignment with Bowtie 2. Nat Methods 9:357–359.

123. Gruber-Vodicka HR, Seah BKB, Pruesse E. 2020. phyloFlash: Rapid Small-Subunit rRNA Profiling and Targeted Assembly from Metagenomes. mSystems 5:e00920–20.

124. Li D, Liu C-M, Luo R, Sadakane K, Lam T-W. 2015. MEGAHIT: an ultra-fast single-node solution for large and complex metagenomics assembly via succinct de Bruijn graph. Bioinformatics 31:1674–1676.

125. Meyer F, *et. al*. 2022. Critical Assessment of Metagenome Interpretation: the second round of challenges. Nat Methods 19:429–440.

126. Wu Y-W, Simmons BA, Singer SW. 2016. MaxBin 2.0: an automated binning algorithm to recover genomes from multiple metagenomic datasets. Bioinformatics 32:605–607.

127. Kang DD, Li F, Kirton E, Thomas A, Egan R, An H, Wang Z. 2019. MetaBAT 2: an adaptive binning algorithm for robust and efficient genome reconstruction from metagenome assemblies. PeerJ 7:e7359.

128. Olm MR, Brown CT, Brooks B, Banfield JF. 2017. dRep: a tool for fast and accurate genomic comparisons that enables improved genome recovery from metagenomes through de-replication. ISME J 11:2864–2868.

129. Chaumeil P-A, Mussig AJ, Hugenholtz P, Parks DH. 2019. GTDB-Tk: a toolkit to classify genomes with the Genome Taxonomy Database. Bioinformatics btz848.

130. Shannon P. 2003. Cytoscape: A Software Environment for Integrated Models of Biomolecular Interaction Networks. Genome Research 13:2498–2504.

131. Brown CT, Olm MR, Thomas BC, Banfield JF. 2016. Measurement of bacterial replication rates in microbial communities. Nat Biotechnol 34:1256–1263.

132. Oksanen J, Kindt R, Legendre P. 2019. vegan: Community Ecology Package. R package version 2.5-6.

